# Optimal metabolic states in cells

**DOI:** 10.1101/483867

**Authors:** Wolfram Liebermeister

## Abstract

Cells, in order to thrive, make efficient use of metabolites, proteins, energy, membrane space, and time. How, for example, should they allocate the available amount of protein to different metabolic pathways or cell functions? To model metabolic behaviour as an economic problem, some flux analysis model, kinetic models, and cell models apply optimality principles. However, due to their different assumptions these models are hard to compare and combine. Benefits and costs of metabolic pathways – e.g. favouring high production fluxes and low metabolite and enzyme cost – can be derived from general fitness objectives such as fast cell growth. To define pathway objectives, we may assume “optimistically” that, given a pathway state, any cell variables outside the pathway will be chosen for maximal fitness. The resulting fitness defines an effective pathway objective as a function of the pathway variables. Here I propose a unified theory that considers kinetic models, describes the set of feasible states as a state manifold and score each state by cost and benefit functions for fluxes, metabolite concentrations, and enzyme levels. To screen the state manifold and to find optimal states, the problem can be projected into flux, metabolite, or enzyme space, where effective cost and benefit functions are used. We reobtain existing modelling approaches such as enzyme cost minimisation or nonlinear versions of Flux Balance Analysis. Due to their common origin, the different approaches share mathematical optimality conditions of the same form. A general theory of optimal metabolic states, as proposed here, provides a logical link between existing modelling approaches and can help justify, interconvert, and combine metabolic optimality problems.

## 1 Introduction

Living cells cannot be understood in all their complexity, and since we cannot understand them, we need to describe them by simplified pictures. Blueprints of cellular networks, depicting biochemical reactions and regulation arrows, can be translated into mathematical models whose variables and equations reflect network structure, chemical dynamics, and molecular regulation. In such models, cells are often portrayed as well-mixed chemical reaction systems (as if they were little test tubes) or information-processing devices (as if they were little computers). These pictures highlight dynamics and regulation and may tell us how cells behave – but now why they behave as they do. In fact, cells are also believed to save resources such as nutrients, proteins, energy, intracellular space, to use opportunities in their ecological niches, and to find compromises between opposing needs. Beyond metabolic networks (the “blueprint”) and mechanistic models (for dynamic simulation), this is a third level of description of cells, one in which how processes or compounds are associated with costs and benefits and in which we ask how they *should* be used [1]. This description of cells does not only concern mechanisms, but functional benefits, selection advantages or, as it were, the “economy of the cell”. In an economic metaphor, we may compare a cell to a company that can produce different goods and makes choices about investments depending on the market (a) Yeast proteome (b) Cost and benefit in a metabolic pathway (c) Benefit terms in enzyme space prices of materials, energy, and the goods themselves. In this analogy, metabolic pathways would correspond to commodity or value chains.

The idea of optimal resource allocation underlies many topical questions in cell biology, and specifically metabolism. For example, what determines the choice between different pathways, e.g. between fermentation or respiration for ATP regeneration, or the preference for specific carbon sources? How should metabolism and protein production be adapted for fast growth or for an optimal use of resources? Since cellular pathways are tightly coupled, even apparently simple choices (e.g. between alternative pathways) may depend on complex dynamics and resource allocation in other places, involving cellular networks with thousands of variables. If our aim is to describe such choices by simple models that zoom in on a small pathway only one type of variables (e.g. fluxes), we need a good theory that relates optimality in these simple models to optimality in the entire cell (or for that matter, optimality in complex cell models of which our model describes only a tiny part). While such a “zooming” can already be complicated in mechanistic models, optimality models pose the extra challenge that cost and benefit functions have to be translated between model approaches), and that any subsystem must be optimised *assuming that* all other subsystems are optimised as well.

In the economy of the cell, metabolic enzymes play a main role. Enzymes steer and enhance metabolic rates, but their production, maintenance, and space requirements also put burdens on cells. The thrust to keep this burden low shapes metabolic fluxes and proteomes. Figure 1 (a) shows typical protein investments in yeast: almost half of the protein budget is devoted to metabolic enzymes, large investments go into glycolysis, and individual enzyme abundances are largely different. This raises questions about dynamics, but also about optimal allocation of protein and metabolite resources. How can protein profiles be explained or predicted? How do they depend on metabolic network structure, on the demand for metabolic fluxes, on enzyme kinetics, and on enzyme sizes? And can the demands for individual enzymes be predicted from models? As external conditions are changing, the cell’s protein investome may be affected in various ways. For example, a shortage of iron may affect the expression of both iron-converting and iron-containing proteins, with indirect effects on other processes and on the entire proteome. What choice of enzyme levels will maximise cell fitness, and what metabolic states emerge from these choices? How can optimal enzyme investments be predicted from network structure, enzyme kinetics, and thermodynamics? And how can we use this knowledge in practice to predict fluxes or to engineer metabolic pathways?

**Figure 1:**
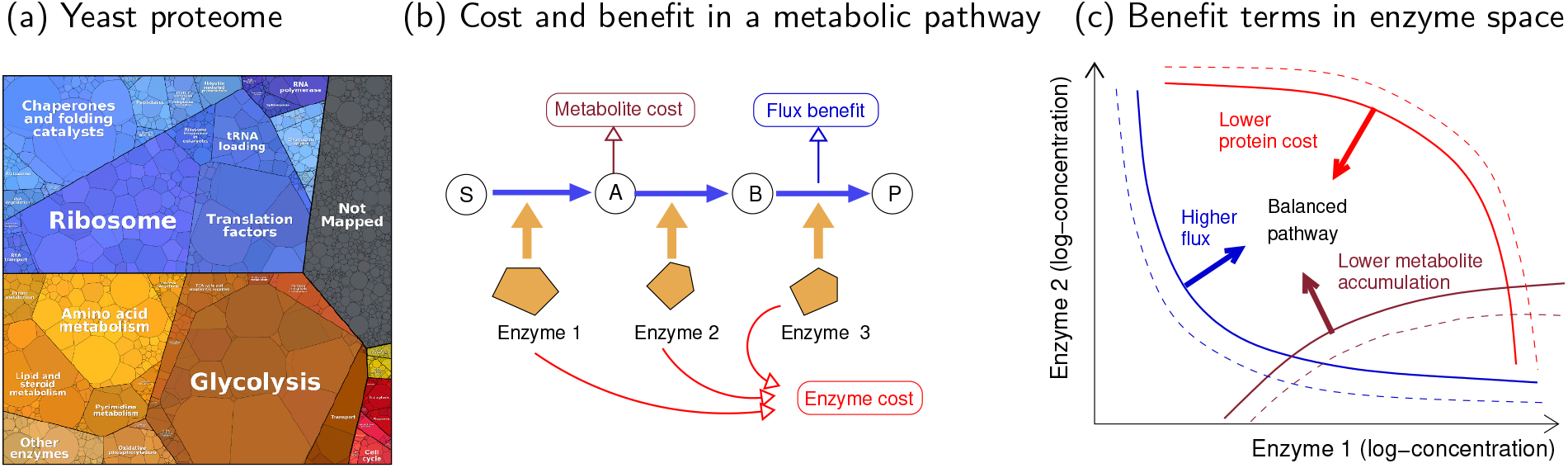
Protein profiles in cells explained by a compromise. (a) Measured protein abundances (molecule count numbers) in the budding yeast *S. cerevisiae* [2] (data from [3]). Proteins are represented by polygons (with sizes representing abundance) and coloured and arranged by cellular functions (larger coloured areas). (b) Optimal enzyme profile in a simple metabolic pathway. Enzyme levels determine the metabolite concentrations **c**^st^ and fluxes **v**^st^ in steady state, as described by a kinetic model. Fluxes are scored by a benefit function *b*(**v**), while metabolite and enzyme levels are scored by cost functions *g*(**c**) and *h*(**e**). (c) Enzyme space. Each possible enzyme profile is represented by a point (only two dimensions are shown). Benefit and cost functions *b*(**v**^st^(**e**)), *g*(**c**^st^(**e**)), and *h*(**e**) shown by contour lines (redrawn from [4], where a similar model has been analysed). Enzyme profiles that allow for a high metabolic flux (blue lines) at low metabolite concentrations (brown lines) and enzyme levels (red lines) provide a selection advantage. If these three target terms are described as benefits (flux benefit, negative metabolite cost, and negative enzyme cost), the sum of benefits is maximal in a point in which the three gradients cancel out (“balanced pathway”). How can we obtain these functions? The (negative) protein cost is a simple function in enzyme space. To obtain the other two gradients, we need to solve the kinetic model for a steady state depending on the enzyme vector **e**.

Protein allocation has been studied by various types of biochemical network models, which describe metabolites, reactions, and their catalysing enzymes. There are two main types of models (see Box 1): kinetic models, which describe the dynamics of metabolite concentrations, enzyme concentrations, and fluxes; and Constraint-Based Models (CBM) such as Flux Balance Analysis (FBA) which assume stationary flux distributions but ignore metabolite concentrations and enzyme kinetics. Aside from shared network structure, different modelling paradigms may also share some equations, e.g. the stationarity condition (setting the net production or consumption of internal metabolites to zero). We can imagine chemical conversion as a flow of matter in the metabolic network. For example, glucose molecules enter the cell and are converted, step by step, into carbon dioxide: we can imagine the carbon atoms “flowing” through the reaction network. Due to this flow, metabolite concentrations can change in time: if the net production rates (production minus consumption) of a metabolite are not zero, the concentration will increase or decrease. A steady state is a state in which all internal metabolite concentrations are mass-balanced, and therefore constant. This puts constraints on the possible flux distributions. Kinetic rate laws describe how reaction rates depend on metabolite concentrations: the higher the substrate concentrations (and the lower the product concentrations), the higher the flux, in line with termodynamic laws. The dynamics of concentrations and fluxes are tightly coupled: metabolite concentrations determine the fluxes through kinetic rate laws, and fluxes determine the changes of metabolite levels by mass balance. Biochemical network models can be coarse-grained or fine-grained and describe smaller or larger parts of the cellular systems. They can cover various cell processes including metabolism, cell signalling, gene expression, protein and mRNA synthesis, or describe an entire cell. On a more abstract level, different types of models can describe either *possible* cell states (e.g. constraint-based flux models), *actual* dynamics (e.g. kinetic metabolic models), or *desired* behaviour (e.g. optimality-based models).

### Box 1

**Metabolic optimality problems**

The modelling approaches below describe different parts of the cell (a metabolic pathway, metabolism as a whole, or systems beyond metabolism) and contain different types of variables (including fluxes, metabolite concentrations, and enzyme levels). In optimality-based models, biological objectives are optimised under physical and physiological constraints. In this article we ask how the different modelling paradigms are logically related.

#### Flux balance analysis

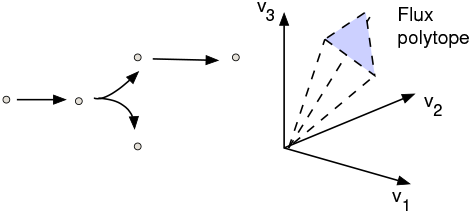

##### Stoichiometric flux-balance models

Flux Balance Analysis (FBA) models (see Eq. (8)), predict metabolic fluxes from network structure, stationarity, and optimality criteria such as high metabolic production (benefit) or low enzyme demand (cost). Based on cost and benefit functions for fluxes, the aim is to find flux distributions with a maximal benefit under flux constraints [5], a maximal benefit at a given flux cost [6], or a minimal cost at a given benefit [7]. Constraint-based models do not describe enzyme kinetics: enzyme levels are either ignored or treated by simplified rules, e.g. assuming that enzyme levels and fluxes are proportional. Linear FCM (i.e. FBA with a minimal weighted sum of fluxes) usually uses heuristic cost weights, representing the enzyme burden per flux in each metabolic reaction. If we assume constant metabolite concentrations, such a weighted flux minimisation is actually equivalent to a minimisation of enzyme burden, and the weights are simply given by 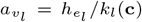 (**c**) (with enzyme cost weights 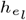, metabolite concentrations *c*_*i*_, and catalytic rates *k*_*l*_(**c**). Metabolite concentrations can be considered, but only to formulate thermodynamic constraints on the flux directions.

#### Kinetic pathway model

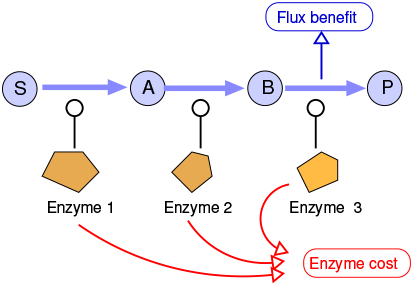

##### Kinetic models with enzyme optimisation

Kinetic models describe the dynamics of compound concentrations. They can be used to model metabolism, signalling, gene expression, or the allocation of protein resources to metabolism or ribosomes. Enzyme levels, fluxes, and metabolite concentrations are linked by rate laws. Given all enzyme levels and initial conditions, a metabolic state can be computed by numerical integration. To find optimal metabolic states, the enzyme levels may be optimised^1^, e.g. to provide large production fluxes at low total enzyme amounts (see Figure 1) [10, 11], i.e. we minimise the fitness Eq. (1) under the constraints Eq. (5). To model metabolism in growing cells, the dilution of compounds can be taken into account. The growth rate can be treated as a given parameter, as a dynamic variable, or as a variable to be optimised.

#### Cost minimisation in metabolite space

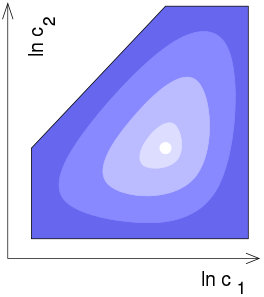

##### Enzyme optimisation at given fluxes

Given a desired metabolic flux distribution, we may search for metabolite and enzyme levels that realise the fluxes at a minimal biological cost. Such enzyme and metabolite concentrations follow from an optimality problem in metabolite space [12], called enzyme cost minimisation (ECM): we consider a kinetic model, choose a flux distribution, and search for enzyme and metabolite concentrations that realise these fluxes at a minimal cost. The cost function may include a direct metabolite cost and a cost for the enzyme levels required. Given metabolite concentrations and fluxes, the enzyme levels are obtained by a simple formula, and the resulting optimisation is a convex optimality problem in log-metabolite concentration space. The resulting cost defines an effective kinetic flux cost function for FCM.

#### Models of growing cells

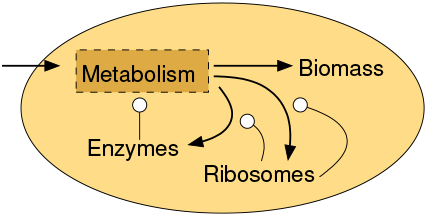

##### Optimal resource allocation for cell growth

In microbial cell models, possible objectives are, for example, maximal cell growth, growth at minimal nutrient levels, or survival at a maximal temperature (requiring expression heat shock proteins). To keep all cell compounds at physiological levels and to compensate for their constant dilution, the cell needs to reproduce all metabolites, enzymes, ribosomes, and so forth. Cell models can be constraint-based or kinetic. Resource balance analysis [13, 14] is a constraint-based method that covers metabolism, enzyme and ribosome production, and macromolecular processes such as protein folding by chaperones. Fluxes and enzyme levels are treated as strictly proportional. The aim is to obtain a steady state at a fixed growth rate, and to determine the maximal growth rate (i.e. dilution rate) at which this condition can be met. Kinetic cell models are usually smaller. The aim may be to maximise growth by optimising the production of different sorts of proteins, for protein production (catalysed by ribosomes) or metabolism (catalysed by metabolic enzymes), [15], the kinetic constants that determine the allocation of protein may be optimised.

To explore the idea of cellular resource allocation, we need to describe different possibilities of a system under constraints (e.g. a given amount of space that can be filled by compounds). To translate this into optimality problems, we consider a mechanistic biochemical model and postulate that some biological objective needs to be optimised, possibly under constraints. Some model variables (e.g. enzyme levels) are treated as “choice variables”, that is, as controllable parameters that are not described mechanistically (e.g. by assuming a biochemical model of enzyme production and degradation), but *chosen* to optimise the objective function. Whenever model parameters (e.g. external concentrations) are changing, the control variables must be adapted to ensure optimality [16].

Understanding (and predicting) optimal metabolic states is hard because metabolic fluxes, metabolite concentrations, and enzyme levels are linked in two ways: on the one hand via physical laws and dynamics (rate laws, stationarity conditions, and physiological bounds), and on the other hand via optimality (assuming that each variable is optimised *given* the other ones). In contrast to purely mechanistic models, our model solution must not only be physically correct, but also economically self-consistent. An optimal state is characterised both by quantitative variables (e.g. optimal fluxes or concentrations) and by qualitative choices, e.g. which pathways are used, which enzymes are suppressed, and so on. In all these choices, the enzyme efficiencies play a key role. If all metabolite concentations were known, we could compute the catalytic rates (or “enzyme efficiencies” defined as the the flux per enzyme level). If these catalytic rates were constant, each reaction would require a fixed enzyme level per flux, i.e. fluxes and enzyme levels would be proportional. The resulting optimality problem can be solved by linear FCM (i.e. FBA with a minimal sum of weighted fluxes). But in fact, in order to know the metabolite concentrations, we already need to know the optimal cell state! So we need to find all variables (fluxes, metabolite concentrations) at once. In reality, all variables are changing: efficiencies and metabolic strategies depend on each other, and our challenge in modelling is to break this circle! In any case, we see: good estimates on enzyme efficiencies can provide us with good approximate solutions. Therefore one of our main aims is to predict enzyme efficiencies in optimal states.

To optimise cell objectives such as fast growth or life at limited nutrients, cells need to behave economically. In individual pathways they must realise compromises between opposing needs, e.g. between the cost and benefits of biochemical processes. Compromises may be governed by a “willingness to pay” principle, stating that a cost in one place is acceptable if it enables an equally high (or higher) benefit elsewhere. This principle can be used to make costs and benefits comparable in the entire cell. Mathematically, compromises in biochemical networks can be modelled by optimality problems: we consider compound concentrations and reaction rates, list all physical and physiological constraints, and optimise an objective such as a minimal use of resources or maximal cell growth. Figure 1 (b) shows an example, the search for optimal enzyme levels in a pathway with three target functions: a benefit for fluxes, a cost for enzyme levels, and a cost for metabolite concentrations. Trade-offs between targets can be modelled in different ways: by combining them into one objective (e.g. the flux benefit per enzyme cost); by optimising one target while which fixing the others; or by multi-objective optimisation [20]. In all these cases, a condition for optimal states is that gradients of the target functions must be balanced (see Fig (c)). This will be discussed below. The optimisation of metabolic states takes places under various kinds of constraints such as physical constraints on state variables (e.g. the stationarity condition; rate laws between enzyme levels, metabolite levels, and fluxes; and bounds on concentrations (imposed by limited space, thermodynamics, or general cell physiology). The constraints determine a set of feasible states. In addition, there can be constraints on metabolic targets: for example, we may minimise the enzyme demand (a metabolic target) at a fixed biomass production rate (another metabolic target). By constraining the targets, e.g. limiting the total enzyme budget, we obtain a feasible region in the state manifold for our optimality problem. Below we consider the minimisation of enzyme cost at a fixed flux benefit as a running example. Starting from Eq. (1), we require a predefined flux benefit and ignore the metabolite cost. Then, we write enzyme cost (originally, a function of enzyme levels) as a function in the space of fluxes and metabolite concentrations.

Metabolic models come in various forms: some are kinetic, others constraint-based; some describe metabolism, others include protein production and cell growth; some use metabolic targets (e.g., maximising a pathway flux), others use whole-cell objectives (e.g. maximising cell growth). These differences make different models hard to compare and combine. Metabolic optimality problems can be grouped in several ways (Table 1). First, by what variables appear in the model, e.g. enzyme levels, metabolite concentrations, and fluxes, and whether they are treated as parameters, control variables, or dependent variables. Second, by how cell variables are scored by metabolic targets, such as ATP production or the sum of absoute fluxes. Third, by how compromises between these targets are described mathematically. To trade benefit against cost, we can maximise their weighted difference, maximise benefit at a fixed cost, minimise cost at a fixed benefit, or perform a multi-objective optimisation. Thus, the same biological problem can be modelled in different ways. In this article, we ask: how are these different formulations of a problem related, and how can they be derived from each other, unified, or combined? To obtain a unified framework, let us see what different modelling approaches have in common. First, they all rely on network models and share the same underlying network structure. Second, different models may describe the same pathways with the same biochemical parameter values. Third, different optimality problems (e.g. optimisation of enzyme levels, or optimisation of metabolite concentrations at given optimal fluxes) turn out to be mathematically equivalent, leading to the same predictions. Generally, if different optimality problems, for the same mechanistic cell model, make the same assumptions (e.g. that high enzyme levels are costly), either explicitly (by a penalty term for high enzyme levels) or implicitly (because enzyme production, described explicitly in the model, consumes resources), will they also yield the same results? And if so, what are the reasons for this equivalence?

**Table 1:** Modelling approaches describing compromises between metabolic cost and benefit. The control variables in optimality problems may be enzyme levels, metabolite concentrations, or fluxes. Metabolic states may be scored by different cost and benefit functions, and trade-offs may be modelled by maximimising a benefit-cost difference, by maximising a benefit at a fixed cost, by minimising a cost at a fixed benefit, or by multi-objective optimisation.

|  | Enzyme levels | Metabolite concentrations | Reaction fluxes | Growth rate |
| --- | --- | --- | --- | --- |
| Maximise benefit<br>(at given cost) | Flux maximisation [17] |  | Molecular crowding<br>FBA [6] | Cell models<br>[15, 14] |
| Minimise cost<br>(at given benefit) | ECM [12] | ECM [12] | FBA with minimal<br>fluxes [7, 18] |  |
| Optimise weighted<br>difference | Optimal enzyme<br>levels [10] | ECM [12]<br>(with benefit term) |  |  |
| Multi-objective<br>optimisation |  |  | Multiple flux<br>objectives [19] |  |

In this article, I consider metabolic optimality problems and describe them by a unified framework from which various metabolic optimality problems, as well as their optimality conditions, can be derived. I start with an overview of existing modelling paradigms. Then I focus on simple optimality problems like in Figure 1 and ask how their optimal states can be characterised. Hence, for a general framework, we consider metabolic states as points in a combined flux/metabolite/enzyme space. The set of feasible states, respecting all laws and constraints of the model, is a high-dimensional manifold, and a search for optimal states can be formulated as an optimality problem on this manifold. Starting from this general problem, we can derive specific optimality problems for fluxes, metabolite levels, or enzyme levels. In these problems, the original fitness function must be replaced by *effective* objectives in the “visible variables”, which implicitly account for the other “hidden” variables. To unify these different problems again, we consider optimality conditions for hypothetical state variations and write them as cost-benefit Compromises between opposing targets may be formulated in different ways (e.g. maximising flux benefit at a fixed enzyme budet, or minimising the enzyme demand of a given flux profile). Despite these different model formulations, the optimality conditions have the same shape: this confirms that the optimality problems themselves are equivalent. The argument will be further extended in [21], where I introduce a local economic theory of cellular networks in which the contributions of enzymes, metabolites, and reactions to cell fitness by economic values, comparable to prices in economics. The values follow general laws that have the form of local balance equations, suggesting that value (in optimal states) is a conserved quantity that flows through the network. The theory holds for general cell models based on biochemical networks.

## 2 Metabolic optimality problems

Depending on the mathematical formalism used, cell models can describe different processes and employ different variables. Here we focus on metabolisc pathways or networks and adopt a framework that covers kinetic and stoichiometric models. The variables – fluxes, metabolite concentrations, and enzyme concentrations (in vectors **v, c**, and **e**) – are interlinked by kinetic rate laws, mass balance, and other constraints (such as bounds on the total molecule concentration in cells). To keep things simple, we focus on steady states and ignore metabolite dilution in growing cells. We consider an optimality problem for a metabolic pathway^2^ and describe a pathway’s contribution to cell fitness by a flux benefit *b*(**v**), a metabolite cost *g*(**c**), and an enzyme cost *h*(**e**), and assume that the cell maximises a fitness function

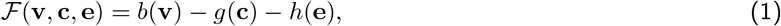

given by a difference of three targets^3^. By convention, fluxes are scored by benefits (to be maximised) while concentrations are scored by costs (to be minimised). The terms in Eq. (1) may also be weighted by prefactors, but here we ignore such prefactors (assuming that the targets are scaled appropriately). Maximising a fitness (1) is not the only way to model trade-offs: alternatively, we may optimise one of the targets while keeping the others fixed or we may apply apply multi-objective optimisation between the three targets. This will be discussed below.

Our fitness function (1) tells us which combinations of fluxes, metabolite concentrations, and enzyme levels would be desirable, but it does not tell us which metabolic states are actually possible. To define physically feasible states, we use a kinetic model that links a flux vector **v**, metabolite profile **c**, and enzyme profile **e** by rate equations and enzymatic rate laws:

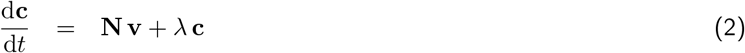

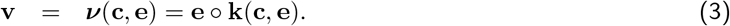

The term *λ* **c** describes dilution in growing cells and is often ignored (assuming that the cell growth rate *λ* is slow compared to typical metabolite turnover times). The symbol ∘ denotes a componentwise vector multiplication. We consider rate laws of the form ν_*l*_ = *e*_*l*_ *k*_*l*_(**c**), with the enzyme level *e*_*l*_ as a prefactor and a catalytic rate *k*_*l*_ depending on metabolite concentrations (also called apparent *k*_cat_ value). Note that the vector **c** contains internal metabolite concentrations (determined by model equations), while “*external* “ metabolites (with fixed concentrations) appear as parameters in the rate laws *k*_*l*_(·)^4^. We assume that all reactions are enzyme-catalysed and that all enzymes are fully specific^5^. Typical rate laws are, e.g. mass-action or Michaelis-Menten kinetics. Here, we allow for any reversible rate laws that are differentiable and thermodynamically consistent (as explained below) [22, 23]. The catalytic rate (assuming a flux in forward direction) can be written as *k*(**c**) = *k*_cat_ *η*^rev^(***θ***) *η*^kin^(**c**), where the turnover rate *k*_cat_ is the maximal rate of reaction events per enzyme molecule and the thermodynamic forces *θ*_*l*_ depend on the metabolite concentrations. From the rate law, we can define the enzyme demand function:

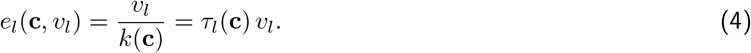

The inverse *τ*_*l*_ = 1*/k*_*l*_, as “reaction slowness”, denotes the flux-specific enzyme demand (mM enzyme per mM/s flux) and, at the same time, the average time between microscopic reaction events for a single enzyme molecule. In stationary states (also called steady states), metabolite concentrations and fluxes are constant in time. If we ignore dilution, the vectors **v**^st^ and **c**^st^ must satisfy

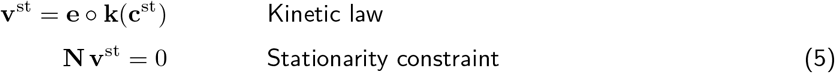

The stoichiometric matrix **N** determines two important features of metabolic models, stationary fluxes and conserved moieties, respectively related to the right-kernel matrix **K** (satisfying **N K** = 0) and the left-kernel matrix **G** (satisfying **G N** = 0). Columns of **K** (and their linear combinations) describe all possible stationary fluxes. Rows of **G** (and their linear combinations) describe conservation relations between internal metabolites (e.g. the total number of phosphate groups in the system, assuming that phosphate groups can be transferred between molecules, but cannot enter or leave the system). If conserved moiety concentrations are given (in a vector **c**_cm_), we can apply the constraint

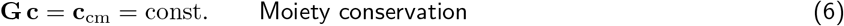

For each column **g** of **G**^T^, the product **g** · **c** is constant in time. In models with moiety conservation, **N** can be split into a product **N** = **L N**_R_ where the rows of the reduced stoichiometric matrix **N**_R_ refer to independent internal metabolites. The link matrix **L** relates rate variations *δ***r**^ind^ of independent metabolites to the resulting variations of all metabolites, *δ***r** = **L** *δ***r**^ind^.

State variables may be restricted by physiological bounds (e.g. minimal and maximal metabolite concentrations, or an upper bound on their sum). Specifically, some models constrain compound concentrations by positivity constraints, lower and upper bounds, or density constraints (on the overall concentration of metabolites, enzymes, or both, in cell compartments). In contrast to conserved moiety constraints, these constraints are usually inequality constraints, e.g.

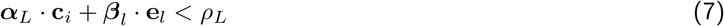

for compartment *L*. A density constraint may be used to reserve space for water (in the cytosol) or lipids (in membranes) or to limit the osmotic pressure.

If we restrict our model to fluxes, require stationary fluxes, and put lower and upper bounds on individual fluxes (e.g. defining flux directions and maximal reaction velocities), we obtain constraints for plausible flux distributions:

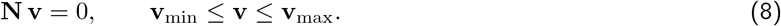

Flux Balance Analysis (FBA) [24] uses these constraints and applies additional optimality criteria to select a single flux distribution. In classical FBA, we maximise a linear benefit function *b*(**v**) corresponding to the first term in Eq. (1). A typical benefit function is the biomass production rate, i.e. the flux in the biomass-producing reaction. In FBA with molecular crowding, we consider a weighted sum of the fluxes as a proxy for enzyme demand and constrain it by an upper bound, assuming **v**^st^ = **e** ∘ **k** and Σ*β*_*l*_ *e*_*l*_ ≤ *ρ*. In Flux Cost Minimisation (FCM), conversely, we fix a flux benefit value (e.g. the biomass production rate) and minimise a flux cost *a*(**v**), for example the sum of absolute fluxes (in the case of FBA with minimal fluxes) or a weighted sum of absolute fluxes as a proxy for enzyme cost. Biologically meaningful flux cost functions should increase with the absolute flux |*v*_*l*_|. Therefore, the costs must show a minimum at *v*_*l*_ = 0 and that the scaled derivatives (∂*a/*∂*v*_*l*_) *v*_*l*_ must be positive^6^.

Our formalism for pathway models, together with density constraints and dilution, can also be used to build whole-cell models. For example, the pathway reactions in Figure 1 may be seen as nutrient import, metabolism (producing precursors), and protein synthesis. The enzymes catalysing the three reactions (transporters, metabolic enzymes, and translation machinery including ribosomes) compete for protein resources, and in the model we may either fix the protein budget, or assume that protein is costly. Mathematically, we can minimise the enzyme demand or penalise enzyme usage by a cost (assuming that enzyme usage in one pathway reduces the protein budget available for other pathways (non-modelled), which would lower the cell’s performance).

Metabolism is shaped by thermodynamic laws. Each reaction has a thermodynamic driving force *θ* = −∆*G*/*RT*, where ∆*G* is the reaction Gibbs free energy, *R* is Boltzmann’s gas constant and *T* is the absolute temperature. Each reaction flux is given by the difference of forward and reverse one-way fluxes *v*_+_ and *v*_−_, with a ratio *v*_+_*/v*_−_ = e^θ^ [26]. For positive forces *θ*, we obtain the formula

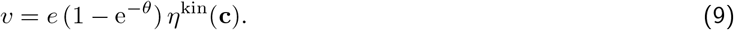

Positive fluxes (where *v*_+_ *> v*_−_) require positive forces while negative fluxes (where *v*_+_ *< v*_−_) require negative forces. In short, fluxes and forces must have the same signs, where zero fluxes are always allowed (“weak sign condition” sign(*v*_*l*_) = sign(*θ*_*l*_) for all *v*_*l*_ = 0). The strong sign condition requires, additionally, that non-zero forces lead to non-zero fluxes: the signs of fluxes and forces are exactly the same, including zero values. The strong sign condition implies that a thermodynamically possible flux cannot be fully suppressed (e.g. by enzyme inhibition or repression).

In a convenient approximation (assuming activity coefficients equal to 1), a reaction’s driving force is a function of substrate and product concentrations. Noting that ∆*G* = ∆*µ* (where *µ*_*i*_ are the chemical potentials of the reactants) and setting 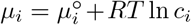 (where concentrations *c*_*i*_ are given in units of a standard concentration), we obtain the formula for driving forces *θ*_*l*_ = −**Δ**_*µl*_/*RT* = ln *K*_eq,*l*_ − Σ*ln*_*il*_ ln *c*_*i*_, where 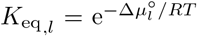 is the reaction’s equilibrium constant. The weak sign condition implies, for all active reactions, a dependence

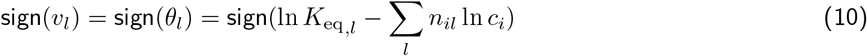

between flux directions and metabolite concentrations: the flux direction in each reaction depends on the ratio of product and substrate concentrations, called mass-action ratio. Below some critical mass-action ratio, called equilibrium constant *K*_eq_, we obtain *θ*_*l*_ *>* 0 and the flux runs in forward direction. Above this value, *θ*_*l*_ *<* 0 and the flux flips its direction. The relation between mass-action ratio, equilibrium constant and flux direction holds for any thermodynamically consistent rate law. This has important consequences for modelling: (i) given metabolic concentrations determine the flux directions; and (ii) given flux directions yield linear constraints on the possible metabolite log-concentrations.

We saw that known flux directions can be used to constrain metabolite concentrations (due to Eq. (10). In addition, metabolite concentrations may be precisely known or physically constrained^8^ (some concentrations may be predefined, and for others upper and lower bounds may be set empirically). The resulting set of feasible metabolite profiles is a polytope in log-metabolite concentration space. If a flux mode contains infeasible loops (which would require chemical potentials to decrease in a circle) [27], this represents a *perpetuum mobile*. Such flux profiles, which cannot be realised by any choice of metabolite profiles, can be discarded. In brief, thermodynamically feasible flux distributions are loopless (and also the opposite holds if no bounds on metabolite concentrations are assumed). If a flux profile leads to an empty M-polytope (no matter if metabolite bounds are considered), it is called *thermodynamically infeasible*. However, flux profiles may also be thermodynamically feasible, but with unphysiological choices of the metabolite concentrations (e.g. extremely high or low concentrations^9^). Such flux distributions, which (together with metabolite bounds) lead to an empty M-polytope are called *thermo-physiologically infeasible*. By restricting metabolite levels to physiological ranges, we can constrain our flux profiles to a set of feasible segments in flux space. Likewise, the thermodynamic relationships between metabolic concentrations, thermodynamic forces, and fluxes, together with the stationary assumption, define a set of feasible states (**v, c**) called the thermodynamic state space 𝒯). It consists of a collection of polytopes in flux/metabolite space, called patches, and states on this manifold can systematically be constructed (see Box 2).

### Box 2

**The thermodynamic state space 𝒯 of a metabolic models**

To characterise feasible metabolic states, we first focus on fluxes and metabolite concentrations. Enzyme levels and enzyme kinetics will be considered later. For now, we just note that any feasible metabolic state can be realised by the right choice of enzyme levels (and upper bounds on enzyme levels can always be satified by a downscaling of enzyme levels and fluxes). In Box 3 we will see how enzyme levels and enzyme kinetics come into play, and that some of the “feasible” fluxe profiles may be excluded because of high enzyme demands.

Hence, a metabolic state will now be described by a flux profile **v** and a metabolite profile ln **c**, the feasible states (**v**, ln **c**) (satisfying stationarity, thermodynamic laws, and physiological bounds) form a set 𝒯. Flux and metabolite profile are related through physical laws. Due to the second law of thermodynamics, the flux directions must agree with the signs of the thermodynamic forces ***θ*** = −**Δ*µ***, where the chemical potentials are given by 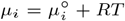 ln *c*_*i*_ (where activitiy coefficients equal to 1 are assumed). In thermodynamically consistent kinetic models, this relationship is automatically satisfied. This relationship between flux and metabolite profiles determines the set 𝒯 of thermodynamically possible states (**v, c**).

We can describe this set by considering the set of feasible force patterns sign(***θ***) = sign(−**Δ*µ***). Each choice of ***µ*** defines possible flux directions, and the states with this pattern form a polytope in *v/m*-space given by the Cartesian product of a flux polytope and a metabolite polytope (see SI) and with box constraints (physiological ranges of fluxes and metabolite concentrations).

Based on the above constraints, feasible polytopes in flux and metabolite space can be constructed. We choose a feasible force pattern sign(***θ***) and consider all compatible flux profiles (or metabolite profiles). For simplicity, we assume that sign(***θ***) does not contain any zero values. If it does, the corresponding fluxes must vanish and the reactant concentrations of these reactions must satisfy linear equality constraints.

1. **Metabolite polytope** Given a force pattern sign(***θ***), we now consider the set of possible metabolite profiles (points in log-metabolite space) compatible with sign(***θ***). Due to thermodynamic constraints (which go hand in hand with thermodynamically feasible rate laws), each metabolite profile determines the flux directions (flux pattern), and so the metabolite space is covered by convex polytopes, each related to one of the flux patterns. The physiological concentration ranges define a feasible box in metabolite space. All M-polytopes outside this box can be discarded. Below we will see that some other polytopes must be discarded too. Altogether, we obtain a collection of feasible M-polytopes, defining the possible metabolite profiles and corresponding flux patterns.

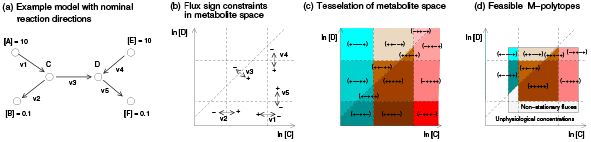
2. **Flux polytope** Each flux pattern belongs to a segment of flux space (i.e. an orthant or a surface of an orthant). Stationarity, and maybe a predefined flux benefit define a feasible subspace in flux space. By intersecting the two, we obtain a feasible flux polytope (S-polytope or B-polytope).

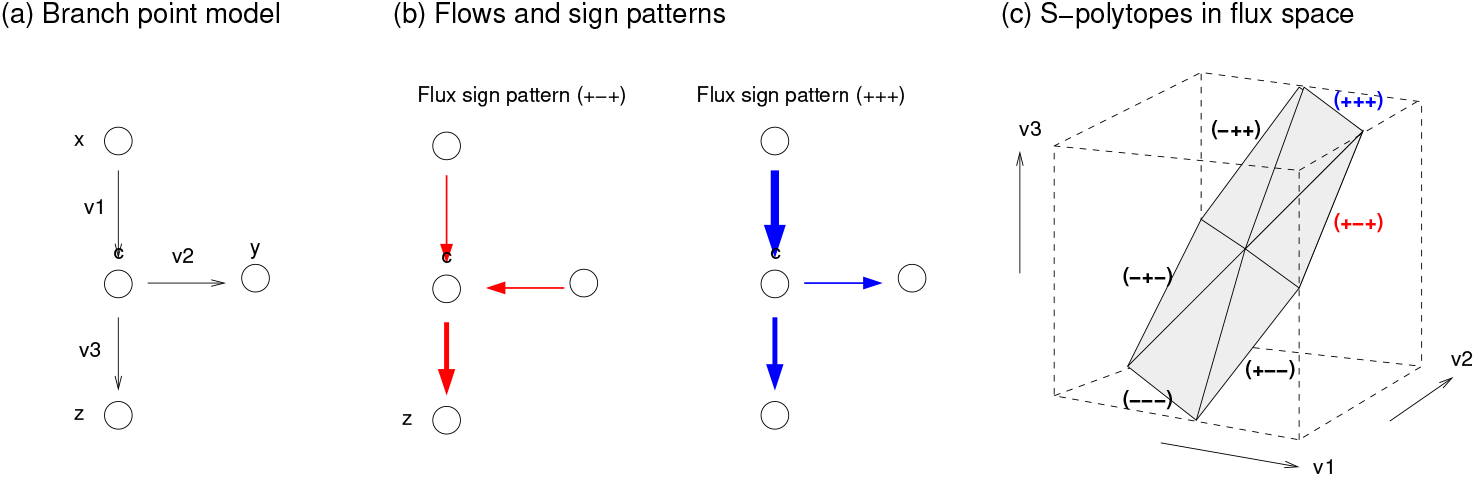
3. Now we can combine the feasible flux and metabolite profiles. For a force pattern to be feasible, there must be non-empty feasible polytopes in flux and metabolite space. If a flux pattern does not allow for a feasible flux profile, the corresponding M-polytope is discarded. If a flux pattern is not realisable by any feasible metabolite profile, the corresponding S-polytope is discarded. By considering all possible force patterns sign(***θ***), we obtain a collection of feasible flux patterns (and corresponding pairs of S-polytopes and M-polytopes) that satisfy all constraints.

I mentioned above that enzyme kinetics is in line with to thermodynamics. First, for realistic models, we need to use reversible, thermodynamically consistent rate laws: the equilibrium constants need to satisfy Wegscheider conditions (that is, they must be derivable from a choice of standard chemical potentials), and equilibrium constants and kinetic constants (*K*_M_ and *k*_cat_ values) in each reaction must satisfy Haldane relationships [28, 22]. Parameter sets that violate these relationships are inconsistent. Second, the energy dissipation per flux (descibed by the unitless driving force *θ*) determines the flux ratio *v*_+_*/v*_−_, and has effects on enzyme efficiencies. Reversible rate laws can be factorised into the terms *v* = *e k*_cat_ *η*^rev^(*θ*) *η*^kin^(**c**) with dimensionless efficiency factors *η*^rev^(*θ*) and *η*^kin^ ranging between 0 and 1 [23]. A low driving force implies a small thermodynamic efficiency *η*^rev^, and therefore a high enzyme demand per flux. In reactions close to thermodynamic equilibrium, there is a trade-off between energy dissipation and enzyme investment: low-yield pathways, which by dissipate more energy, may show a lower enzyme demand.

In our rate laws *v* = *e k*(**c**), the enzyme levels appear as prefactors. If the metabolite concentrations are fixed, enzyme levels and fluxes will be proportional. Hence, we can scale all enzyme levels and fluxes by the same factor, while leaving the metabolite concentrations unchanged (assuming that all reactions are enzyme-catalysed). For example, we can always increase all fluxes until our enzymes hit a density constraint: then, an upper bound on the enzyme levels will define a maximal scaling for fluxes. Knowing this, we can compare flux distributions by their benefit per enzyme amount, which depends only on their shapes, and not on their scaling (and we can compare flux profiles at a unit benefit, assuming that they can always be scaled if necessary).

## 3 The metabolic state manifold

Before we consider optimality problems, we first need to see between which metabolic states, i.e. which arrangements of fluxes, metabolite concentrations, and enzyme levels a cell can choose. The relationships between different variables, including thermodynamics, are summarised in Figure 2. The metabolic variables are coupled by rate laws, stationarity, and thermodynamic laws and may be constrained to physiological ranges. Dependencies between fluxes, metabolite concentrations, and enzyme levels can be described in different ways (see Figure 3), and in metabolic optimality problems the basic variables can be enzyme levels (as in Figure 1), fluxes (as in FBA), or metabolite levels (as in Enzyme Cost Minimisation [12]). To unify these problems, we avoid the distinction between free and dependent variables and consider all variables in one state space. In this space, the manifold of *feasible steady states* – satisfying all physical and physiological constrains – is the set on which we search for optimal states.

**Figure 2:**
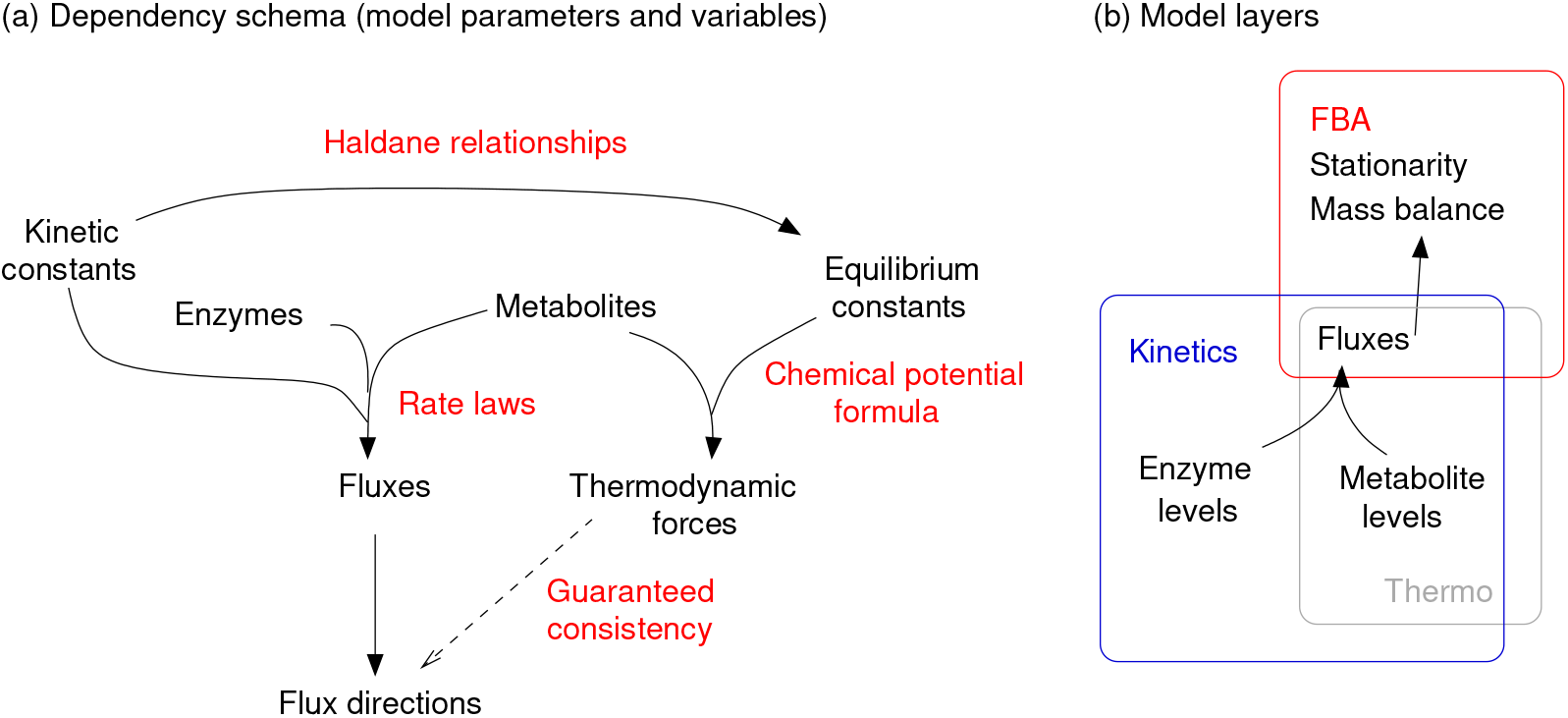
Metabolic variables and layered models. (a) Dependencies between parameters and variables in kinetic metabolic models. Dependencies are shown by arrows (towards dependent variables). The schema shows one possible way to describe the dependencies and to compute quantities from each other (see Fig. 3 for alternatives). If we use thermodynamically correct rate laws and our standard formula for thermodynamic forces (without activity coefficients), the resulting flux directions are guaranteed to follow the thermodynamic forces. In flux analysis, this consistency does not appear automatically, but needs to be guaranteed by extra constraints. (b) A model can be divided into “layers” containing different types of variables and requiring different model approaches. The layers shown correspond to FBA methods (red box), kinetic models (blue box), and reaction thermodynamics (grey box).

**Figure 3:**
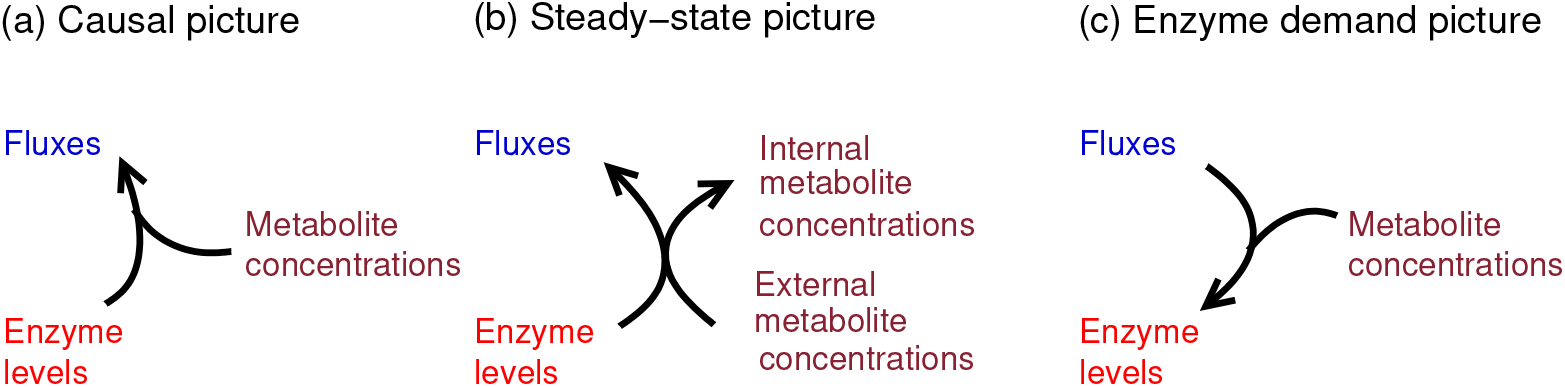
Fluxes, metabolite concentrations, and enzyme levels are linked by kinetic rate laws. (a) In a causal picture, enzyme and metabolite concentrations determine the (possibly non-stationary) reaction rates. (b) In a steady-state picture, the enzyme levels (together with external metabolite and conserved moiety concentrations) determine metabolite concentrations and fluxes. In some cases, the solutions may be non-unique (multistationarity). (c) In the enzyme demand picture, used in enzyme cost minimisation [12], the enzyme demand depends on desired metabolite concentrations and fluxes.

The states of a metabolic system, respecting rate laws, stationarity, and physiological bounds, form a manifold in (**v, m, e**)-space. How can we describe its shape? The manifold *K* is the set of feasible states (**v, m, e**), i.e. points that satisfy all model constraints. The manifold does not imply any notion of independent or dependent variables, but it can be parameterised by **v** and **m**, treating these variables as independent and **e** as dependent. However, not all combinations of **v** and **m** are viable. Combinations that violate thermodynamic laws (and would yield negative enzyme levels if Eq. (4) is applied blindly) must be discarded: thus, all points (**v, m**) must lie in the thermodynamic set 𝒯. Figure 4 shows a simple example, a reversible reaction with a fixed product concentration. In this case, the possible metabolic states form a two-dimensional manifold 𝒦 in (*v, m, e*)-space. Its projection onto the (*v, m*)-plane, the “shadow” 𝒯, consists of two “patches”: one patch (red) for states with high substrate concentrations and positive fluxes, and another one (blue) for states with low substrate concentrations and negative fluxes. One patch corresponds to one flux direction and describes the feasible ranges of metabolite concentrations and fluxes: by taking the Cartesian product of the two ranges, we obtain the patch in flux/metabolite space. States outside the patches (shown in light grey) are thermodynamically infeasible. The state manifold is a curved surface in (*v, m, e*)-space. It consists of two “sheets”, which can be obtained by “lifting” the patches in *e*-direction: for each point (**v**, *m*), the enzyme level *e* is obtained from the enzyme demand function 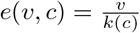, where *k*(*c*) is the catalytic rate of the enzyme. Due to our rate laws, the enzyme demand function has a typical shape (see Figure 4): at a given metablite concentration *m*, the enzyme demand *e* is linear in fluxes; and at fixed fluxes, the demand is convex in log metabolite concentrations *m*. Close to thermodynymic equilibrium, it goes to infinity. Hence, the shape of the manifold is easy to understand. The boundary between the patches depends on the equilibrium constant, while the shape of the manifold above them depends on the rate law: the enzyme level scales linearly with *v* (straight diagonal lines) while the contour lines (intersection lines with *c* − v planes) represent the rate law *v* = *e k*(*c*) for a fixed value of *e*. The state manifold has some convenient properties: it is differentiable and path-connected, i.e. the cell can move between all states on the manifold by smoothly changing its metabolite and enzyme levels. Finally, it is ease to construct because given a flux/concentration pair (*v, m*), the enzyme demand *e* is easy to compute and guaranteed to be positive (because in a feasible patch, our rate law yields a rate *r* that matches the sign of the flux *v*).

**Figure 4:**
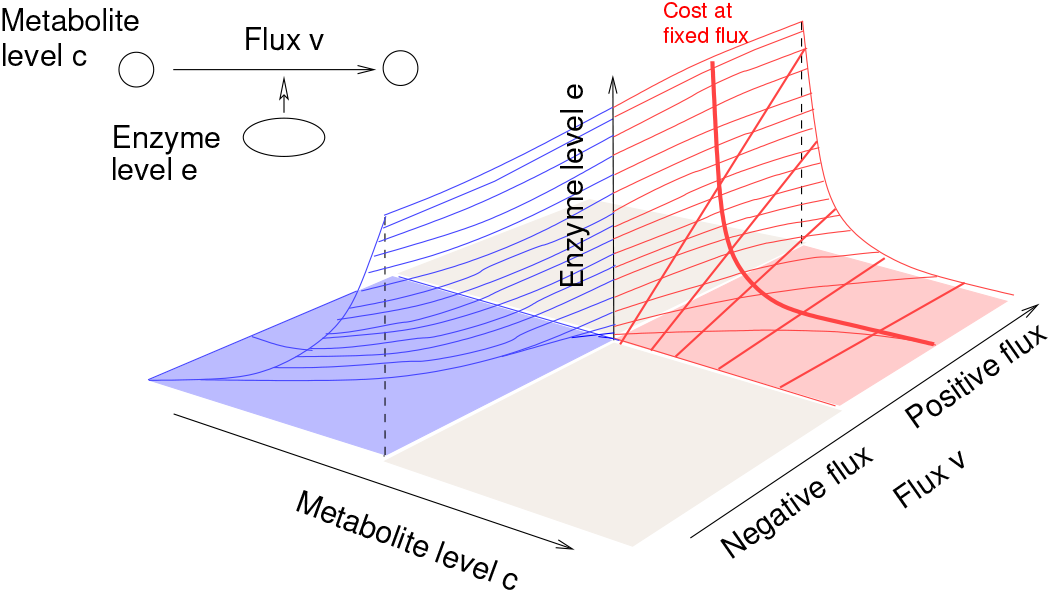
Metabolic state manifold of a single enzymatic reaction with substrate concentration *c*, enzyme concentration *e*, and reaction rate *v* = *e k*(ln **c**). In the example, the product concentration is assumed to be constant. The two patches in (*v, m*) space, corresponding to the two possible flux directions and shown in blue and red, and to sheets of the manifold in (*v, m, e*)-space. Curved contour lines show the rate law at different enzyme levels. Straight lines illustrate that enzyme levels and fluxes are proportional, at a fixed metabolite concentration.

We saw that the two patches touch in one point. In (*v, m*)-space, our cell needs to go through this state (the chemical equilibrium state where *v* = 0) to flip the flux direction. In the state manifold, this point corresponds to a semi-infinite line between the two sheets (with zero flux, equilibrium concentration, and arbitrary enzyme levels). This means: in the chemical equilibrium state, with a zero flux, the enzyme demand *e*(*v, c*) is underdetermined. To enforce a unique solution, we may postulate that cells repress unnecessary enzymes, and set the enzyme levels to *e*_*l*_ = 0 (“principle of dispensable enzymes”).. This yields a bijective mapping between states (**v, m**) and (**v, m, e**), and therefore between 𝒯 and 𝒦. However, as we can see from Fig. (4), the manifold 𝒦 will not be continuous anymore, but interrupted by an empty line between the two sheets.

In general, the state manifold is a multidimensional algebraic manifold^10^. While it may have a complicated shape, it can be constructed in a simple way^11^ (Box 3) and has some convenient mathematical properties. (i) A model with *n*_*r*_ enzyme-catalysed reactions and *n*_*ext*_ external metabolites has an *n*_*r*_ + *n*_*ext*_-dimensional manifold which is continuous in almost all points. (ii) The manifold is path-connected, i.e. the system can move between any two points without leaving the manifold^12^ (at least if the system may pass through the equilibrium state **v** = 0). Extra constraints, for example a given flux benefit, lead to a lower-dimensional manifold that may consist of separate regions. (iii) The manifold is differentiable in all points (since we can parameterise the manifold by **c** and **e**, and the rate laws ν_*l*_(**c, e**) are differentiable). Details about state manifolds are given in SI section **??**.

### Box 3

**The kinetic state manifold 𝒦 of metabolic models**

In kinetic metabolic models, the set of feasible states (**v, m e**) is an algebraic manifold called kinetic state manifold 𝒦. It consists of all states (**v, m, e**) that satisfy our model constraints: (i) rate laws for all reactions, defining a mapping (**v, m**) → **e**; (ii) stationary fluxes; (iii) predefined conserved moiety concentrations. We require that rate laws must be thermodynamically consistent, i.e. they must show a zero flux if the mass-action ratio matches the equilibrium constant and positive (negative) fluxes below (above)! Moreover, fluxes and metabolite concentrations may be bounded by physiological ranges, and metabolite and enzyme concentrations must be non-negative. How is this manifold structured? Each state (**v**, ln **c**) defines the required enzyme levels via the enzyme demand function (4).

To construct the manifold as a whole, we first think about possible flux patterns: each state (**v, m, e**) has a flux pattern, and each feasible flux pattern allows for a set of solutions (**v, m**), and therefore for solutions (**v, m, e**). To be feasible, a flux pattern must be realisable by stationary fluxes (possibly, with a predefined flux benefit) and by metabolite profiles that respect physiological bounds and thermodynamic constraints. In brief, a feasible pair (**v, m**) must belong to the thermodynamic state space 𝒯 (see Box 2). Moreover, for each pair (**v, m**) in 𝒯, there is exactly one enzyme profile **e** (given by the enzyme demand function) that leads to a feasible state (**v, m, e**).

The sets 𝒦 and 𝒯 are related via a simple mapping: by applying the enzyme demand function **e**(**v, c**), each patch in 𝒯 can be “lifted” to yield a sheet of the manifold 𝒦. Conversely, by ignoring **e**, each sheet of 𝒦 can be projected again to a patch in (**v, m**)-space. Each patch belongs to a specific flux pattern and is given by the Cartesian product of a feasible flux polytope (in flux space) and a feasible metabolite polytope (in log-metabolite space).

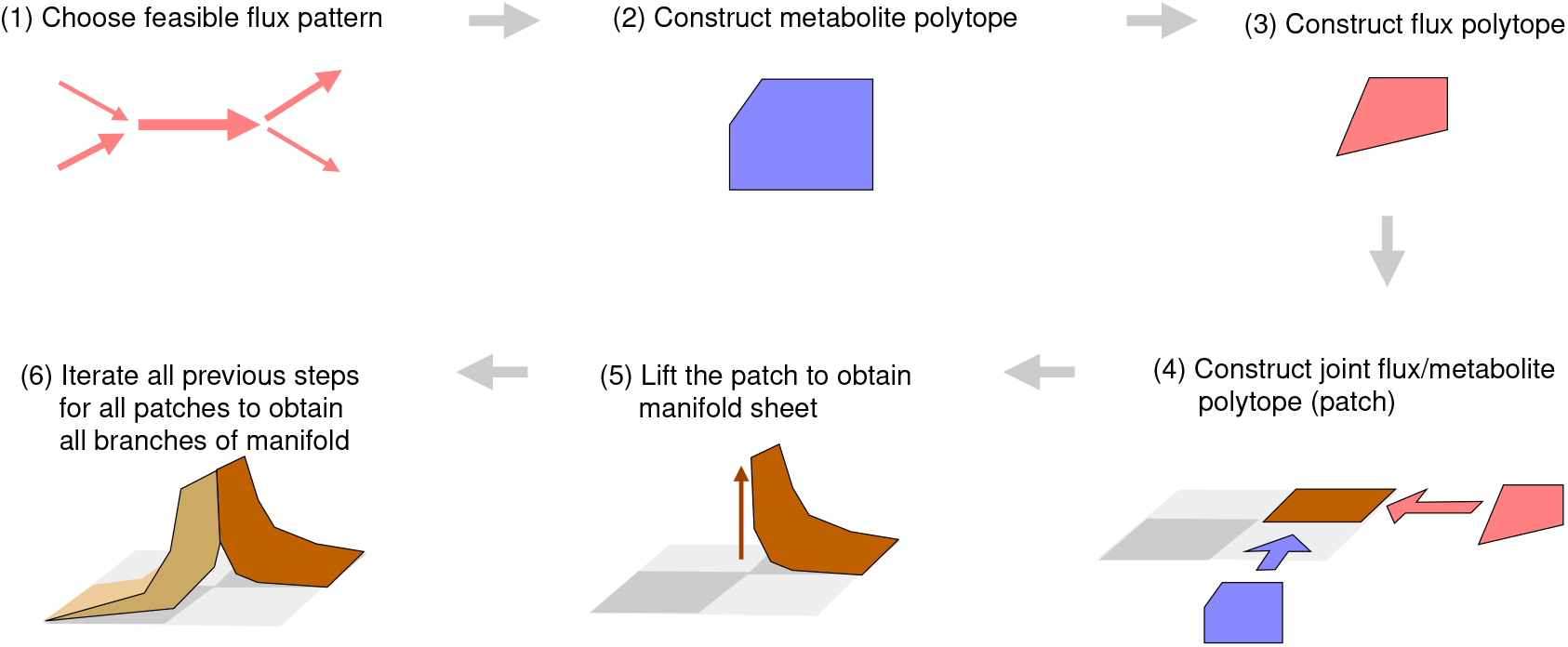

The entire state manifold for a given kinetic model can be constructed following the same logic (see graphics). We first determine the set of possible flux patterns (1). For each flux pattern, we construct the feasible flux polytope (2) and metabolite polytope (3). The Cartesian product of the two polytopes yields a feasible patch (4), like in Figure 4. Each point (**v, m**) from this patch can be realised with exactly one easily computable enzyme profile (the only exception being states with vanishing fluxes), which can be easily computed with the demand function (3). By doing so, we lift each patch (5) to a sheet in (**v, m, e**)-space. All sheets together form the state manifold (6).

We saw that, by generating all feasible flux and metabolite profiles and solving for the enzyme demand, all possible states can be constructed. By enumerating the flux patterns, constructing the flux-metabolite patches, and lifting them using the function *e*(*v, c*), we can systematically construct the state manifold. Our construction of the state manifold shows how closely thermodynamic and kinetic models of metabolism are related. A feasible thermodynamic state (**v**, ln **c**) (a point of 𝒯) must satisfy the relation sign(**v**) = sign(***θ***(ln **c**)), while a feasible kinetic state (**v**, ln **c, e**) (a point of the 𝒦-manifold) must satisfy the kinetic rate laws (3). With thermodynamically consistent rate laws, the second condition implies the first: for each kinetically feasible state (**v**, ln **c, e**), the pair (**v**, ln **c**) will be thermodynamically feasible. Likewise, each feasible thermodynamic state (**v**, ln **c**) can be completed uniquely by an enzyme profile **e** to obtain a kinetically feasible state (**v**, ln **c, e**). We obtain a one-to-one mapping between thermodynamic and kinetic states, with only one exception: if a reaction rate vanishes, this may be due to a vanishing enzyme level, a complete inhibition, or chemical equilibrium. In the latter two cases, the catalysing enzyme can have any concentration: **e** is non-unique. Except for this case, the “thermodynamic” and “kinetic” descriptions are completely equivalent: two models with different rate laws have the same state manifolds, just “distorted” in *e*-direction: there is a state manifold mapping across models (of the same network, but with different rate laws). All this means that thermodynamic flux analysis (e.g. in thermodynamic FBA) is a perfect basis for kinetic modelling: all thermodynamically feasible states, and only those, can be kinetically realised, and exactly in one way! In the thermodynamic picture, a state is just feasible or infeasible. In the kinetic picture, states “close to being infeasible” become enzymatically costly: if we approach a chemical equilibrium state while maintaining a constant flux, the lower and lower driving force must be compensated by larger and larger enzyme levels [12]. In the equilibrium state itself, the enzyme demand would go to infinity.

Importantly, an optimality problem for fluxes, metabolite log-concentrations, and enzyme levels cannot be convex (proof in section **??**). This holds for any objective function (which we discuss in the following section), because it follows from the very shape of the state manifold. To obtain convex optimality problems instead, we may predefine either the fluxes, metabolite concentrations, or enzyme levels, and optimise over the remaining variables. With fixed fluxes **v**, for example, the optimality problem for **c** (and **e**) becomes convex for a wide range of rate laws [12]. Likewise, at fixed concentrations **c**, enzyme levels and fluxes are proportional, leading to convex problems such as linear FCM or FBA with molecular crowding. As discussed below, a combination of these two methods (fixing either fluxes or metabolite concentrations) allows us to find an optimal state (**v**, ln **c, e**) by a combination of convex and concave optimisation.

When dealing with metabolic steady states, there are often the questions whether enzyme levels (and conserved moiety concentrations) determines a state uniquely (i.e. metabolite concentrations and fluxes) and whether a state is dynamically stable. Regarding uniqueness, even kinetic constants, enzyme levels, and external concentrations are fixed, a kinetic model can have multiple steady states. In our manifold picure, this would mean that the manifold, when projected onto e-space, casts a “multiple shadow”: the mapping *e* → (*v, c*) is not unique, but has multiple function branches. Among the algebraic manifolds, the state manifold has a special property : it can be parameterised by pairs (**v, m**), and be described by a function **e**(**v, m**) on the feasible set 𝒯 in (**v, m**)-space). This means: in “**e**-direction” there exists a simple lifting from 𝒯 to 𝒦 and a unique projection in reverse. However, from and to **e**-space, such a unique projection may not always exist! For example, if enzyme concentrations **e** and external metabolite concentrations **s** are treated as basic variables and **c**^st^ and **v**^st^ as functions of them, these “functions” may not be unique^13^: in this case, the projection of 𝒦 to enzyme space is not injective, and a “lifting” from enzyme space to 𝒦 is not unique. The second question concerns dynamic stability. If a steady state is unstable, then once the system deviates from this state (e.g. driven by chemical noise), the dynamics will drive it farther away. In practical modelling, such states are almost irrelevant, because a system under chemical noise would not stay in such states for a long time, just like a pencil, when put on its tip, will not keep standing like this in reality. Starting from an initial state, the dynamical system will either approach one of the stable steady states or show non-steady (oscillating, diverging, or chaotic) behaviour. Some of the states obtained by our screening may be dynamically unstable. In modelling, it makes sense to detect such states based on the system’s Jacobian matrix, and to discard them.

Let us briefly come back to dilution. During balanced growth, fluxes and metabolite concentrations are coupled in two ways: by rate laws and by the stationarity condition **N v** − λ **c** = 0. Because of the latter condition, a feasible flux profile **v** completely determines the metabolite profile **c**, which requries a different description of the state manifold: flux and metabolite profiles cannot be screened independently anymore. The state space 𝒯 is no longer a collection of simple patches (Cartesian products of F-polytopes and M-polytopes), but is the solution set of **N v** − λ **c** = 0 (a plane in (**v, c**)-space, which yields a curved manifold in (**v**, ln **c**) space), intersected with the set of feasible M-polytopes in **m**-space (for thermodynamic correctness). As before, the state manifold 𝒦 is obtained by “lifting” the state space 𝒯, I will come back to dilution in [21].

## 4 Objective functions in flux and metabolite space

We saw that all feasile metabolic states (**v, c, e**) in a kinetic metabolic model can be constructed by screening the feasible combinations of metabolite concentrations and fluxes. Now we come back to our initial question, the search for optimal metabolic states. We consider a kinetic model (including physical laws and physiological constraints), construct its state manifold, choose an objective function on the manifold (e.g. enzyme plus metabolite cost), and search for optimal states.

In theory, optimal metabolic states can be found by screening all feasible combinations of stationary fluxes, metabolite concentrations, and enzyme levels (i.e. all points on the manifold) and choosing the state (**v, m, e**) with the highest objective value. An example is shown in Figure 5. In practice, it may be convenient to hide some of the variables, i.e. to optimise either enzyme levels, metabolite concentrations, or fluxes, but not all of them simultaneously. For this projected optimality problem, we need effective objective functions that describe our objective as a function of the “visible” variables: for example, by optimising an effective objective in flux space, we may obtain the same optimal fluxes we would obtain by optimising over the state manifold. Among the effective objectives, there are two main types: “restricted” objectives, in which the hidden model variables are fixed, and “optimistic” objectives, in which they are assumed to be optimised given the visible variables. By combining the two optimisation procedures, we obtain “layered” models, that is, nested optimisation procedures that combine an optimisation over metabolite profiles and over flux profiles. These procedures are not only useful for numerical optimisation, but also for obtaining optimality conditions for cost and benefit gradients. In [21], we come back to a formulation in which all variables are treated on an equal footing and are linked by explicit constraints.

**Figure 5:**
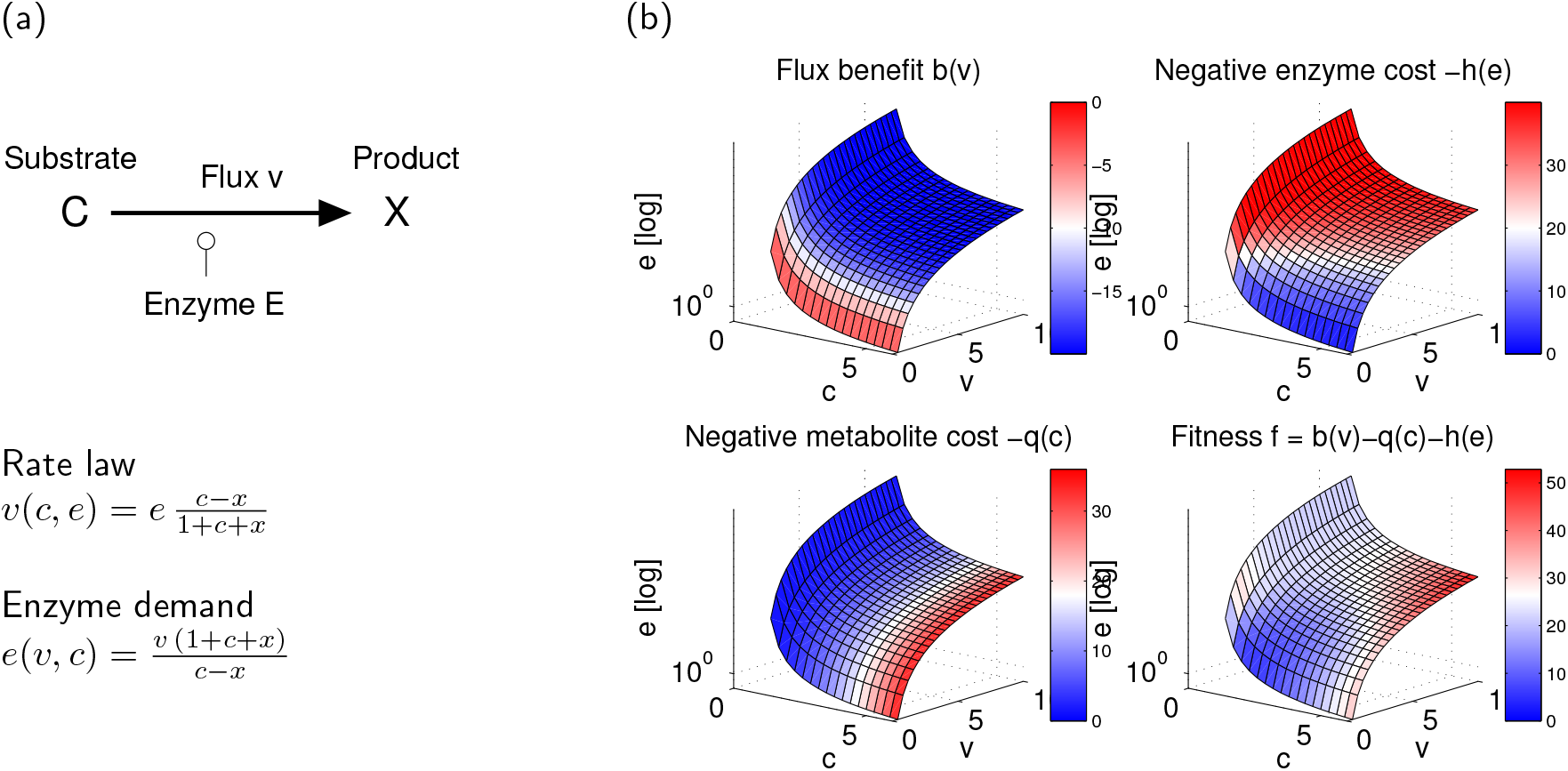
Metabolic states and metabolic objectives in a single reaction (compare Figure 4). (a) We consider a reaction C ↔ X with enzyme level *e*, substrate level *c*, fixed product level *x*, and a reversible Michaelis-Menten rate law (all kinetic constants set to 1 for simplicity). From the rate law ν(*c, e*) we obtain the enzyme demand function *e*(*v, c*). (b) By screening metabolite concentrations and fluxes and computing the enzyme demand, we obtain the set of all feasible metabolic states (*c, v, e*) (similar to Figure 4, but with enzyme level *e* shown on logarithmic scale). In panel 1, the flux benefit function on this manifold (an increasing function of the flux, in arbitrary units) is shown in colours (low: red; high: blue). Panels 2 and 3 show enzyme and metabolite cost (as benefit functions, with a minus sign). The total fitness, in panel 4 (flux benefit minus concentration costs), has a maximum in the center of the blue region.

In metabolic modelling, what matters is often not the entire triple (**v, c, e**), but only a subset of variables, for instance the fluxes. To “hide” the other variables, we project the optimality problem from our state manifold into flux space (see Figure 6): the results is an FBA-like problem in which metabolite and enzyme levels are “hidden” and fluxes are the only model variables. But what about the fact that flux profiles require different enzyme or metabolite concentrations, which makes them differently costly? In general, variables (fluxes, metabolite concentrations, or enzyme levels) can be eliminated in different ways: we can (i) treat some variables as dependent (e.g. write enzyme levels as a function of fluxes and metabolite concentrations), (ii) treat some variables as given (e.g. the metabolite concentrations), or (iii) assume that the hidden variables will always be chosen optimally (given the visible variables). For example, we assume that in the triple (**v, m, e**), the enzyme levels should be hidden and fluxes and metabolite concentrations should remain visible. To hide the enzyme levels, we can (i) set all of them to predefined values, (ii) compute them from metabolite concentrations and fluxes via the rate laws (see Figure 3c), or (iii) assume that all fluxes are realised by cost-optimal enzyme levels. By combining options (ii) and (iii), we can hide metabolite and enzyme levels and obtain an optimality problem only for fluxes. However, “secretly”, each flux profile **v** also defines optimal metabolite and enzyme levels and thus a (minimal) enzyme cost.

**Figure 6:**
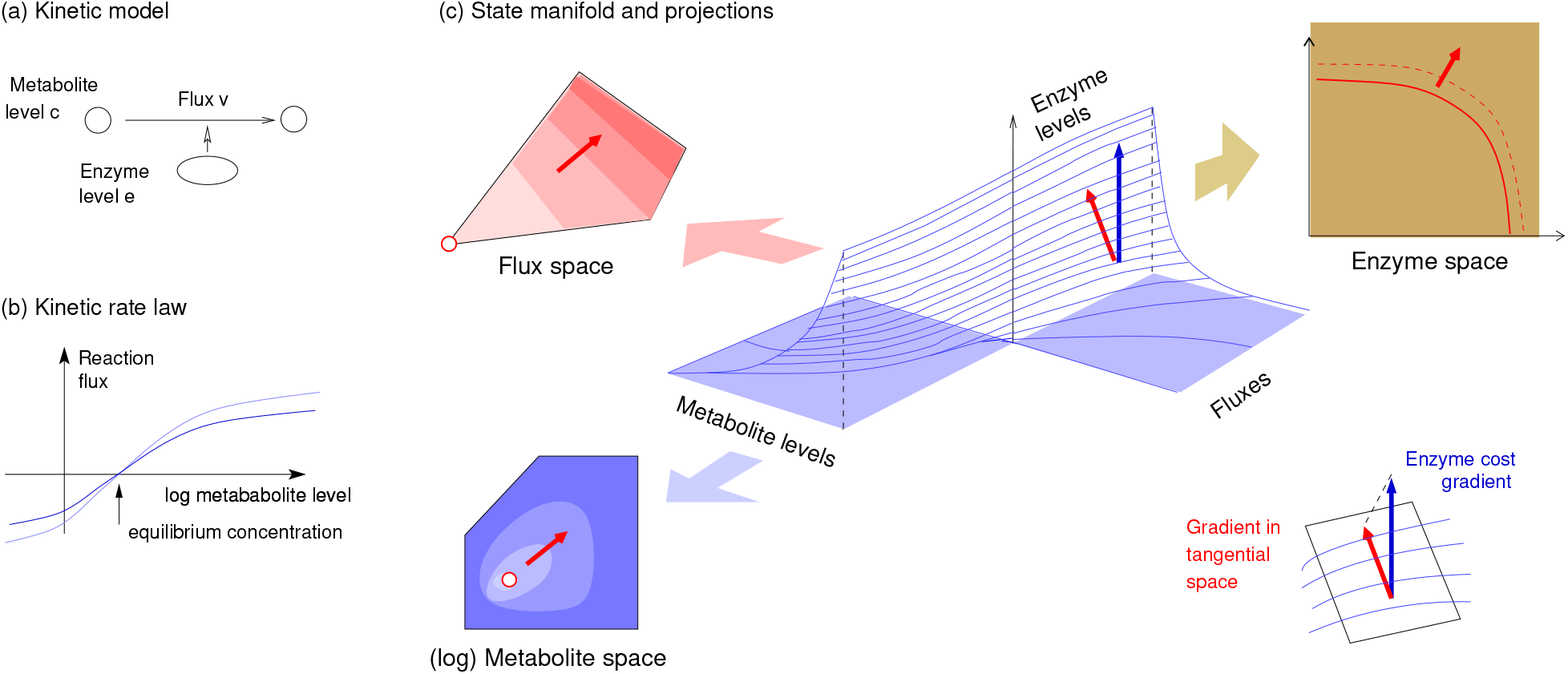
From metabolic model to metabolic optimality problems. (a) Example model: a single reaction. (b) The reaction rate depends on substrate concentration and enzyme level. (c) Metabolic state manifold and optimality problems in flux, metabolite, or enzyme space. Optimality problem on the state manifold (center) and projections into enzyme, flux, and metabolite space. An enzyme cost function (blue gradient shown) defines a gradient in the (tangential space of the) manifold (red). A projection to different subspaces leads to different optimality problems.

As an example, we consider a linear enzyme cost function *h*(**e**) on the state manifold. To hide the enzyme levels (i.e. to remove them from the free variables), we write them as a function of metabolite concentrations and fluxes via the enzyme demand function^14^ **e**(**c, v**). We obtain a number of effective enzymatic cost functions in metabolite (M) and flux (F) space, with convenient mathematical properties (see Table 2 (a) and compare Figure 7).

**Table 2:**
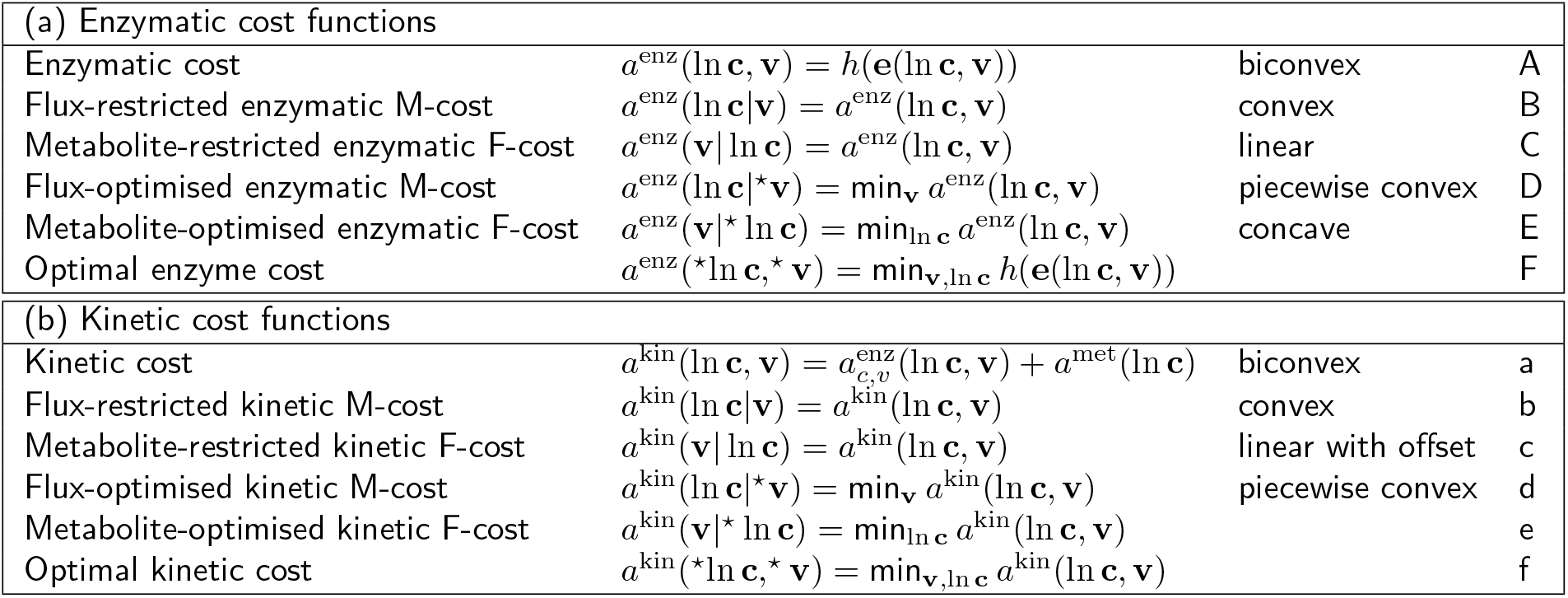
Cost functions in metabolite (M) and flux (F) space. (a) Enzymatic cost functions. The last column refers to the letters in Figure 7. Variables marked by a star ^⋆^ are assumed to be optimised given the visible variables (the resulting objective function is called an “optimistic objective”). All functions are continuous and, except for the flux-optimised M-cost function, differentiable. The enzymatic cost function is biconvex in ln **c** and **v** (but not convex, and linear in **v**). While the cost function *h*(**e**) is a direct cost (i.e. it is define in this way in the model), the enzymatic F-cost *a*^enz^(**v**| ln **c**), for example, is an overhead cost: it is not directly caused by the fluxes, but only implied (i.e. it cannot be avoided given these fluxes). (b) Kinetic cost functions. The functions resemble the enzymatic functions in the table above, with a few differences: for instance, the metabolite-restricted kinetic F-cost has a concentration-dependent positive offset, and the metabolite-optimised kinetic F-cost may be non-concave.

**Figure 7:**
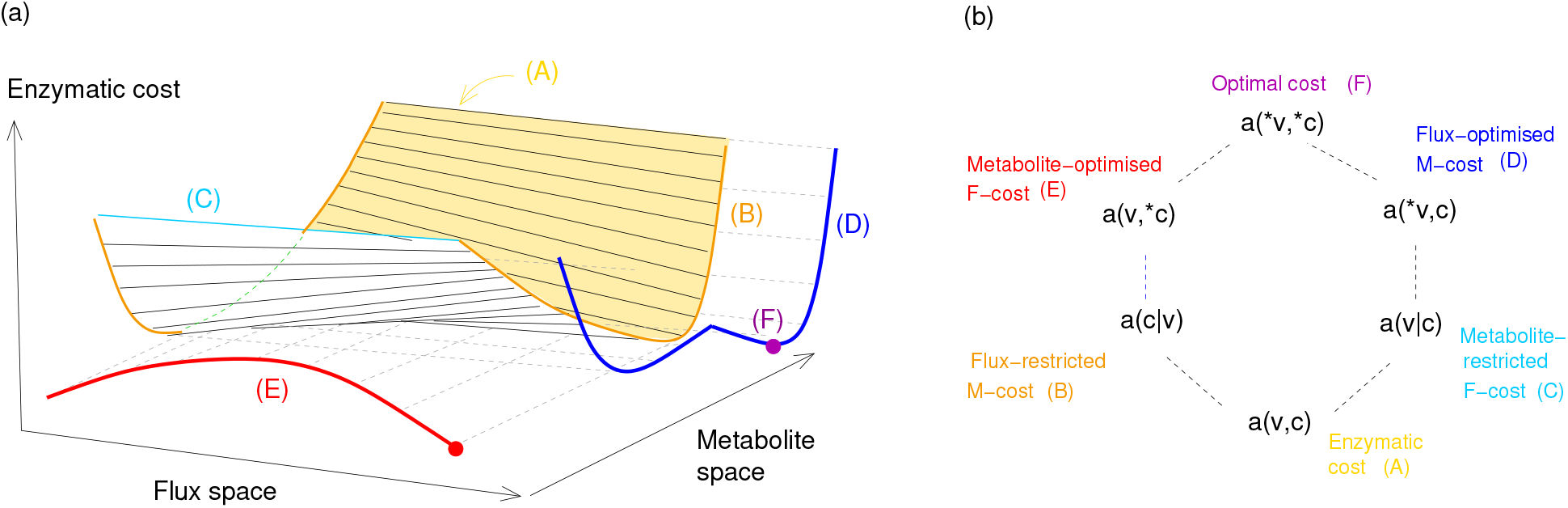
Enzyme cost function in flux and metabolite space. (a) Enzyme cost (hypothetical model, not shown) as a function *h*(**e**(**c, v**)) of fluxes and metabolite concentrations (yellow surface). From this function we obtain effective (“restricted” or “optimistic”) enzyme cost functions in flux or metabolite space (horizontal axis). (b) Relations between the cost functions from (a). Based on the enzymatic cost *h*(**e**(**c, v**)) (bottom), restricted and optimistic cost functions are obtained by constraining or optimising some of the variables.

Projections can be applied to any objective function on the state manifold (i.e. any function of fluxes, metabolite concentrations, and enzyme levels). For example, the kinetic cost function *a*^kin^(ln **c, v**) can be defined by considering a convex metabolite cost *a*^met^(ln **c**) together with our enzyme cost (representing, e.g., the total cell volume occupied by metabolites and enzymes). In the different subspaces, we obtain the effective objectives shown in Table 2 (b).

## 5 Optimal metabolic states

Having learned about state manifolds and objective functions, we now come back to optimality problems in which we evaluate benefit and cost terms (e.g. the ones from Eq. (1)) while restricting the state variables **v, c**, and **e** to feasible states (see Figure 5). We already know (from Section 3) that such optimality problems cannot be convex. To optimise on the state manifold, we need to be able to screen it systematically, and we already know how to do this. We know this already. Instead of varying fluxes, metabolite concentrations, and enzyme levels independently (which would violate the constraints and make us leave the manifold), we consider a complete set of basic variables, e.g. fluxes and metabolite concentrations, and write our optimality problems in these variables. In this way, our optimality problem can be projected from the state manifold to flux, metabolite, or enzyme space (see Figure 6), with effective objectives given in the tables above. Starting from simple objective functions (like the terms from Eq. (1)) and treating them in flux, metabolite, or enzyme space leads some well-known metabolic optimality problems (see Figure 1 and Box 1). Let us see a few cases.

In our first optimality problem, kinetic models with optimal enzyme levels, we predefine the conserved moiety concentrations, treat and enzyme levels as control variables, and determine steady-state metabolite concentrations **c**^st^(**e**) and fluxes **v**^st^(**e**). With the steady-state enzyme benefit function *q*(**e**) = *b*(**v**^st^(**e**)) − *g*(**c**^st^(**e**)), we write the fitness Eq. (1) as a function of **e** alone:

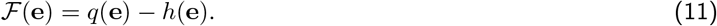

Since the functions **c**^st^(**e**) and **v**^st^(**e**) are not available in closed form and need to be evaluated numerically^15^. Since the fitness landscape ℱ(**e**) may have multiple local optima, optimisation is numerically demanding. Luckily, there is an alternative: with fluxes and metabolite concentrations as control variables and enzyme levels as dependent variables, the fitness can be written as

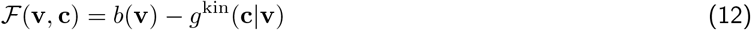

with an effective cost function *g*^kin^(**c**|**v**) = *g*(**c**) + *h*(**e**(**v, c**)). The function ℱ(**v, c**) can be optimised with respect to **v** and **c**. For simplicity, we assume a predefined flux benefit 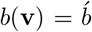 and minimise the enzyme cost at this flux benefit as described above. To perform this optimisation, we combined a minimisation in flux space and metabolite space [25]:

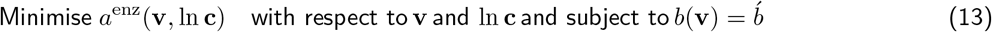

The optimisation with respect to **v** and ln **c**, as a nested optimisation, can be done in two ways (Figure 8).

**Figure 8:**
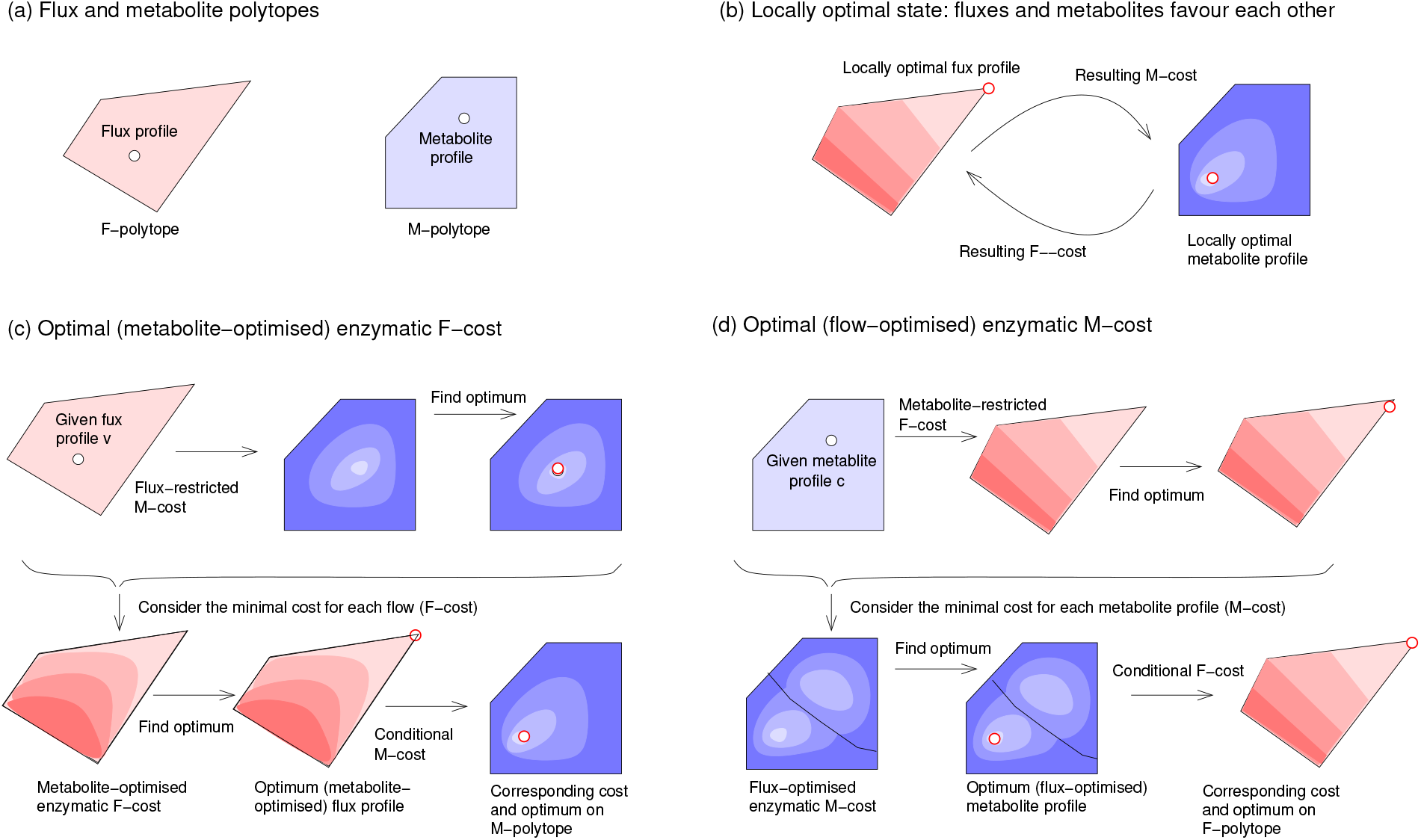
Finding fluxes and metabolite concentrations that minimise enzyme cost. (a) Flux polytope and metabolite polytope for given flux directions (schematic). Fluxes **v** need to be stationary and provide a predefined linear benefit, while metabolite log-concentrations *m* must be thermodynamically consistent with the flux directions. (b) Local optimality condition for **v** and **m**: the flux profile must be optimal given the metabolite profile and vice versa. (c) Minimising the metabolite-optimised enzymatic F-cost *a*^enz^(*v*| ⋆ *c*). For each flux profile **v**, the metabolite log-concentrations are optimised for a minimal cost (top). The resulting enzyme cost function (formally, an optimistic F-cost) is a concave function on the flux polytope (bottom). (d) Minimising the flux-optimised enzymatic M-cost. For each metabolite profile **m**, fluxes are optimised for a minimal cost (top). The resulting optimistic metabolite cost function is piecewise convex on the flux polytope (bottom).

In the first method, we optimise over all metabolite profiles and, for each of these profiles, over the flux profiles (given the metabolite profile). In the inner optimisation (for fluxes), each flux profile **v** has an enzyme demand **e**(**v**|**c**) and the enzyme cost *a*^enz^(**v**|**c**) = **e**(**v, c**), called *metabolite-conditioned enzymatic F-cost*, depends linearly of the fluxes. Minimising this cost in flux space (at a fixed flux benefit) is a simple LP problem (in fact, an FBA with a minimal weighted sum of fluxes). However, this cost still depends on the metabolite profile **c**. As a function of **c**, it can be seen as a cost function *g*^enz*/*v^(**c**) on the metabolite polytope, called flux-optimised enzymatic M-cost. This function is generally piecewise convex, i.e. continuous and convex in regions of the metabolite polytope (but need not be generally convex). It can optimised with respect to **c** (see Figure 7).

In the second method, we optimise fluxes and metabolite concentrations in the opposite order: we optimise over the flux profiles, and for each flux profile we internally optimise over the metabolite profiles. In the inner optimisation (for metabolite profiles), we assume a flux profile **v** and obtain the (flux profile-restriced) enzymatic cost *g*^enz^(**c**|**v**), a cost function in metabolite space which is convex and which we optimise over the metabolite profiles **c**.The resulting cost function, a function of fluxes, is called the *metabolite-optimised enzymatic F-cost a*^enz^(**v**|⋆ **c**) in flux space. Instead of the enzymatic cost function in Eq. (13), we can also consider the kinetic cost, which includes metabolite costs and is obtained by projecting the sum of enzyme and metabolite costs into metabolite space.

A locally optimal metabolic state is a state that performs better than (or equally well as) all similar feasible states. How can we find such states? Let us consider enzyme cost minimisation as an example: to find an optimum, we parameterise the state manifold by fluxes and logarithmic metabolite concentrations *m* and minimise the enzymatic cost *g*^enz^(**m, v**) = *h*(**e**(**c, v**)) as a function of **v** and **m** at a given flux benefit *b*(**v**) = *b*. Since *g*^kin^(**m, v**) is biconvex (see Figure 8), each flux profile **v** defines a unique optimal metabolite profile **m**, and each metabolite profile **m** defines a unique optimal flux distribution **v**. In a locally optimal state, both fluxes and metabolite concentrations must be optimal, that is, they must be optimally adapted to each other. Since we can easily compute the optimal flux profile for a given metabolite profile (by linear FCM), and we can easily compute the optimal metabolite profile for a given flux profile (by ECM). Locally optimla states are easy to find. Since the flux solutions of linear FCM are vertices of the flux polytope, this must hold for all locally optimal metabolic states.

Finally, starting from general optimality problems on the state manifold, how can we derive different versions of FBA? For thermodynamic FBA; we omit the enzyme levels as variables, that is, project the state manifold to (v,c)-space. The projection yields exactly the solution space of thermodynamic FBA, and if the orginal problem considered is “minimising enzyme cost at a given flux benefit”, then also after the projection we can constrain the flux benefit and minimise and (approximate) enzyme cost in flux space. for example, a linearised form of the enzyme cost function.

Finally, we can ask how the state manifold depends on model parameters and what effects this has on their optimal state. The shape of the state space 𝒯 depends on equilibrium constants, external metabolite concentrations, and on the ranges of fluxes and internal metabolite concentrations. The shape of the 𝒦 manifold (the “lifted” version of 𝒯) depends additionally on *k*_cat_ and *K*_M_ values. A change of the equilibrium constant *K*_eq_ will shift the borders of patches in metabolite space. A proportional change of the (forward and backward) *k*_cat_ values would scale the manifold in *e*-direction, while changes of Michaelis-Menten constants would change the shape of the enzyme demand function in other ways.

## 6 Optimality conditions and economic laws

How can optimal state be characterised, i.e. by which general optimality conditions? We saw that optimal flux profiles, as points in the flux polytope, are typically polytope corners. In contrast, if inactive reactions are ignored, metabolite and enzyme profiles are internal optima in their respective spaces. As shown in Figure 1, such internal optima are characterised by a balance of fitness derivatives. On the state manifold, the derivatives can be seen as gradients in the tangential space, but in a projection to flux/metabolite space, the enzyme costs must be expressed as a function of metabolite concentrations and fluxes. Finding the optimal state with different objectives or constraints can be difficult, but the envelope theorem (Box 4) can help us. What are the optimality conditions for metabolic states and what can we learn from them about compromises between different cell variables? To obtain general economic laws for metabolism, we consider again our main optimality problems, cost minimisation in flux/metabolite space and benefit-cost optimisation in enzyme space.

### Box 4

**Optimistic objective functions and the envelope theorem**

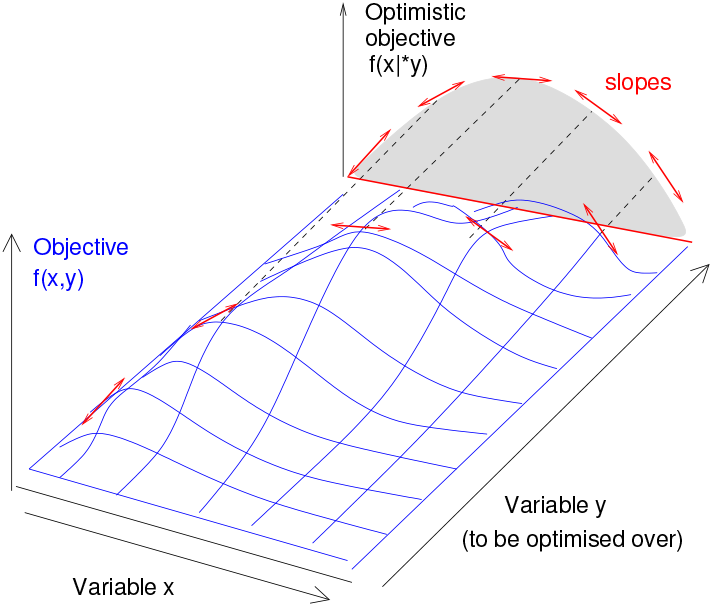

In optimality problem with multiple control variables, we can define an “optimistic objective” for each subset of control variables. In the optimistic objective, the other variables do not appear as function arguments, but are assumed to be optimised. For example, in the image on the left, the projection (or “maximum envelope”) of a function *f* (*x, y*), projected in *y*-direction yields the optimistic function *f* (*x*|^⋆^*y*). The *x*-derivatives of *f* (*x, y*) in the peaks are equal to the derivatives of the optimistic function *f* (*x*,^⋆^*y*).

The enzymatic flux cost *a*^enz^(**v**) = max_**c**_ *h*(**e**) with the constraint **v** = ν(**e, c**) is another example: it follows from the enzyme cost *h*(**e**) by applying the rate laws *v*_*l*_ = *e*_*l*_ *k*_*l*_(**c**) and assuming an optimal metabolite profile **c**. We can also write it as *h*(**v**|**e**,^⋆^*v*). While optimistic objectives may be difficult to compute, their gradients are easily obtained by the “envelope theorem”^16^, a chain rule for optimistic objectives. Let us consider this theorem for a problem with only two variables.

Given an objective function *f* (*x, y*), the y-optimistic objective is defined as *f* (^⋆^*y*) = max_*y*_ *f* (*x, y*). The differentiation rule 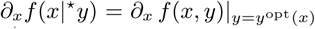 (called envelope theorem) states that the derivative of an optimistic objective *f* ^opt^ is equal to the partial derivative of the objective *f*, taken at the optimal value of *y*. For a minimisation objective *f* (*x, y*) with scalar variables *x* ∈ 𝒳 and *y* ∈ 𝒴, the *y*-optimised optimistic objective is defined as *f* (*x*|^⋆^*y*) = min_*y*∈𝒴_ *f* (*x, y*). The envelope theorem (SI section **??**) states that

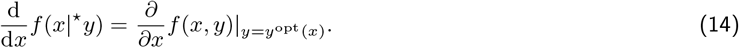

where *y*^opt^(*x*) = argmin_*y*_ *f* (*x, y*). The same rule also applies to functions in high-dimensional spaces, for example to functions on our state manifold. For the metabolite-optimised enzymatic F-cost, we obtain the formula

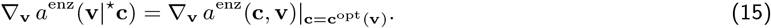

It shows that the gradient of the metabolite-optimised F-cost in a point **v** is equal to the gradient of the metabolite-constrained F-cost, computed at the optimal metabolite profile for flux profile **v**. Similarly, for the flux-optimised enzymatic M-cost the envelope theorem

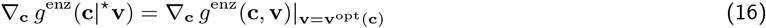

tells us that the gradient of the flux-optimised enzymatic M-cost is given by the gradient of the enzymatic M-cost at the optimal metabolic flux profile for the metabolite profile **c**.

1. **Cost minimisation in metabolite space: metabolite balance condition** To derive our first economic law, we consider cost minimisation in metabolite space (i.e. the ECM problem): given a flux profile **v**, we search for the metabolite and enzyme levels that minimise cost. With metabolite concentrations as the free variables [12], the optimality condition for metabolite concentrations **c** with enzyme demand function reads **e**(**c, v**),

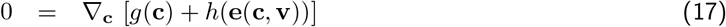

and implies a relation

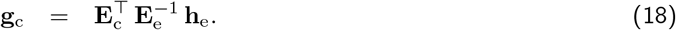

between the cost gradients **g**_c_ (for metabolites) and **h**_e_ (for enzymes) in optimal states^17^ (see Figure 9 (a) and SI section **??**). This *metabolite balance condition* establishes a relation between metabolite prices (in vector **g**_c_) and enzyme prices (in vector **h**_e_). In this vectori equation, each component describes a metabolite and the enzymes that catalyse the surrounding reactions (i.e. those that produce or consume the metabolite, or are regulated by it). Metabolite and enzyme prices are linked by metabolite elasticities 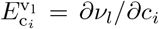 and enzyme elasticities 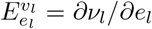. If Eq. (18) holds for ECM problems (with predefined fluxes), it must also hold for FCM problems (where fluxes are optimised along with metabolite concentrations). Eq. (18) has a simple meaning: in an optimal state, a cell cannot improve its fitness by a simultaneous metabolite and enzyme variation that keeps the fluxes unchanged. To see this, we consider a variation *δ***c** of metabolite concentrations^18^. The variation *δ***c** would change the reaction rates by *δ***v** = **E**_c_ *δ***c**, but we compensate this effect by applying a simultaneous enzyme variation. We first imagine an *equivalent* enzyme variation 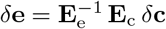, which would have exactly the same effect on fluxes as *δ***c**. If we compensate our perturbation *δ***c** by −*δ***e**, the fluxes remain completely unchanged. This means: any resulting fitness change would have to be entirely caused by the costs of *δ***c** and −*δ***e**, which are given by **g**_c_ *δ***c** and 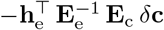. Now we apply the variational principle: in an optimal metabolic state, feasible state variations must be fitness-neutral^19^. Therefore, our compensated variation leave the fitness unchanged 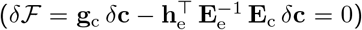. Since this holds for *any* metabolite variation *δ***c**, we can omit *δ***c** from this formula and obtain Eq. (18). Interestingly, all terms in formula (18) describe local quantities (i.e. prices of single metabolites or enzymes and quantitative connections between neighbour variables). However, in models with moiety conservation, this is not the case, beacuse whole sets of metabolite concentrations can be dependent.

**Figure 9:**
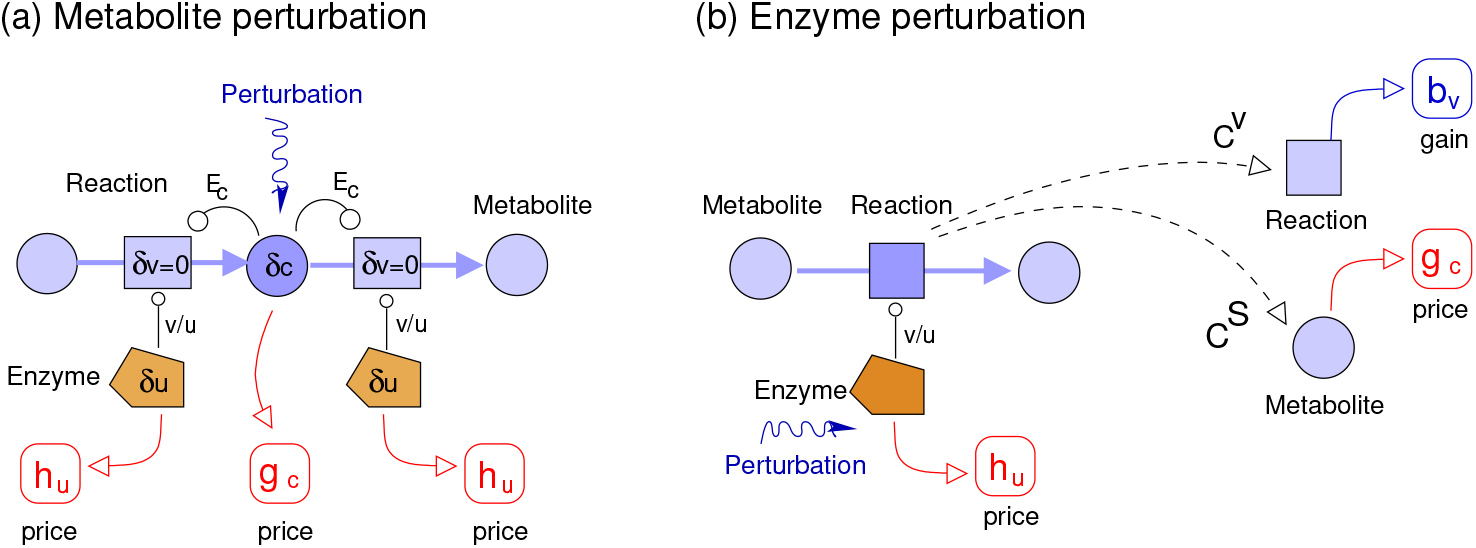
Optimality conditions for kinetic metabolic models. (a) Metabolite balance condition Eq. (18), obtained from cost minimisation at given fluxes. A metabolite variation *δc* with a compensating enzyme variation 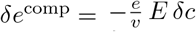 leaves all the rest of the state unchanged. In an optimal state, all such compensated variations must be fitness-neutral (i.e. their cost 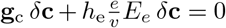 must be zero). In addition, metabolite concentrations can be interlinked via conserved moieties (not shown). (b) Reaction balance Eq. (22), obtained from fitness optimization in enzyme space. A local enzyme variation *δe*, together with the resulting steady-state variations *δv*^st^ and *δc*^st^ (in the entire network), must be fitness-neutral, that is, it must show a vanishing cost: 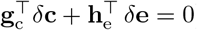).
2. Similarly, an optimisation with objective function (1), with fluxes as control variables, and with metabolite concentrations and enzyme levels as (optimised) dependent variables yields the condition

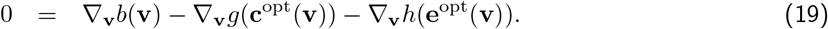

we obtain the balance equation (proof see **??**)

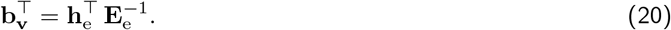
3. **Optimality condition in enzyme space: reaction balance condition** A version of this law for systems with moiety conservation, and a second economic law for flux gains, can be derived from optimisation in enzyme space. We maximise a steady-state Eq. (1), a function of flux benefit *b*(*v*^st^(**e**)), metabolite cost *g*(*c*^st^(**e**)), and enzyme cost^20^ *h*(**e**). The optimality condition for our enzyme profile **e**

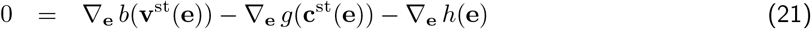

states that the gradients ∇_**e**_ *b*, −∇_**e**_ *g*, and −∇_**e**_ *h* must cancel each other as suggested in Figure 1 (c). By using the theorems of metabolic control theory, we can obtain from this the conditions (proof see SI section **??**)

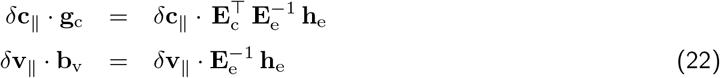

where *δ***c**_||_ or *δ***v**_||_ denote, respectively, legal variations^21^ of concentrations (satisfying moiety conservation) or fluxes (satisfying stationarity). The first equation generalises our economic law (18) to models with moiety conservation (a condition that we had previously not required). For convenience, Eq. (22) can also be rewritten in “point form”. By defining scaled variations (e.g. 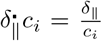), economic variables (e.g. 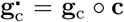), and elasticities (e.g. **Ê**c = Dg(**v**)^−1^ **E**_c_ Dg(**c**)), we obtain the simple laws

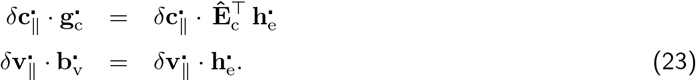

A practical hurdle in using the economic laws in Eqs (22)-(23) is that the elasticities depend on the optimal state and are often unknown.

## 7 Compromises between metabolic targets

Let us now come back to our starting point, metabolic states representing an optimal compromise. The aim of maximising production rates is typically in conflict with (or limited by) nutrient consumption, bounds on specific fluxes, or bounds on some concentrations (e.g. due to limited space, osmolarity etc). Due to compromises in many places, constraints on one variable can have limiting effects on all the others. If different metabolic targets are in conflict, a compromise needs to be found. To describe this, in Eq. (1) we assumed that an optimal state maximises a benefit-cost difference, where cost and benefit terms are functions of model variables such as flux benefit and enzyme cost.

But this is not the only possible way to describe compromises mathematically. In Eq. (1), the metabolic targets are combined into a single objective^22^. Generally, fitness functions may be defined as a difference of flux benefit, metabolite cost, and enzyme cost, but we may consider weighted differences or ratios (e.g. minimise the total enzyme cost per total flux benefit in a pathway). Alternatively, one target may be optimised while keeping the others fixed: e.g. we may maximise flux benefit at a fixed enzyme cost or minimise enzyme cost at a fixed flux benefit^23^. Third, in multi-objective optimisation, we treat the targets as separate objectives and consider Pareto-optimal solutions, forming the Pareto front. A metabolic state is Pareto-optimal if none of the objectives can be improved without compromising the others (in other words, if no other state scores better in one objective and equally well in all others). To construct points on the Pareto front, one may optimise linear combinations of the different objectives (e.g. with randomly chosen prefactors. Alternatively, one may also screen the values of *n* − 1 objectives and, for each choice, optimise the remaining objective.

No matter how we describe our trade-offs, we always obtain the same optimality conditions: the gradients of the target functions need to be balanced! or more precisely, if we write the targets as functions of our choice variables (for example, the enzyme levels) there must be a vanishing linear combination of the gradients (with positive prefactors for cost terms and negative prefactors for benefit terms). Geometrically, this looks exactly like in Figure 1 (c), no matter how we describe compromises (see Figure 10): (i) If we maximise a (weighted) sum of the target functions, the (weighted) sum of their gradients must vanish. (ii) If we optimise one target while keeping the others fixed, in the optimal state a weighted sum of the gradients needs to vanish, with weights given by 1 (for the target to be optimised) and by shadow values for the other targets. (iii) Also in multi-objective optimisation, a weighted sum of the gradients has to vanish (with different weights in different Pareto-optimal states) [20] (proof in SI **??**). Thus, whether metabolic targets are combined into one fitness function, are treated as constraints, or traded against each other in a multi-objective problem, we always obtain the same optimality criterion: a weighted sum of the benefit gradients must vanish! In this sum, metabolic targets that appear as a cost or an upper bound come with a minus sign, while that appear as a benefit or a lower bound with a plus sign. Since all these optimality problems lead to the same optimality conditions, they can be seen as equivalent! The balance of gradients holds for any number of objectives or constraints, e.g. when considering cost terms for different enzyme fractions, in different cell compartments or membranes.

**Figure 10:**
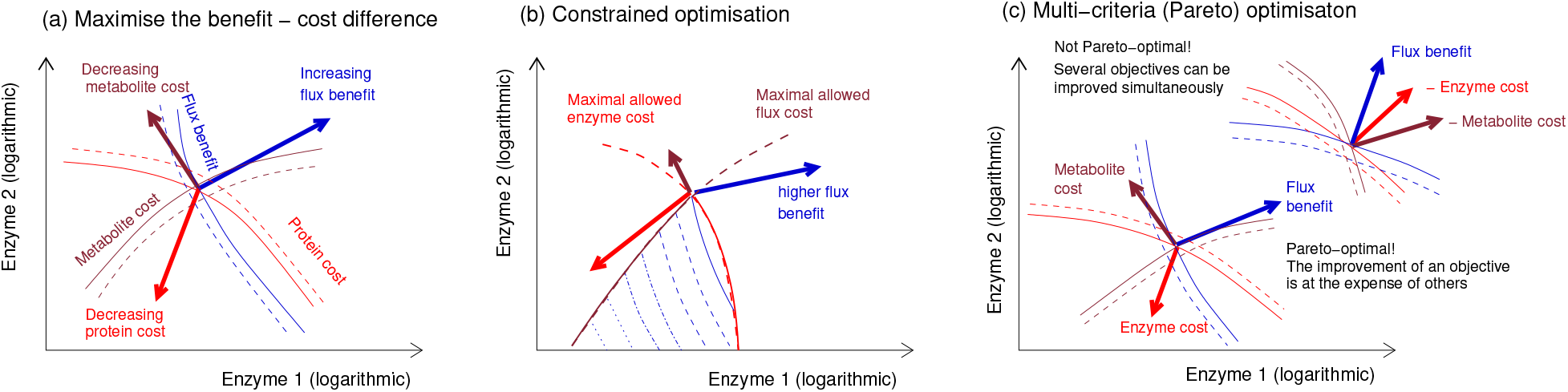
Optimal metabolic states with conflicting targets. Schematic view of an optimality problem in enzyme space (compare Figure 4) with three target functions: flux benefit, metabolite cost (negative metabolite benefit), and enzyme cost (negative enzyme benefit). The panels show different ways to describe compromise. (a) Combined objective function. We assume a (weighted) sum of all benefit terms as the objective. In the optimal state, the three (weighted) gradients must cancel out. (b) Constrained optimisation. Here we maximise flux benefit, while some minimum metabolite and enzyme benefits are required. Again, a weighted sum of the gradients must vanish in optimal states. Some prefactors (the ones that arise from constraints) are given by Lagrange multipliers. (c) Multi-objective optimisation. For an optimal compromise (Pareto-optimal state), again the gradients must be balanced (i.e. some weighted sum needsto vanish). In a non-optimal state (upper right), the gradients point in similar directions and no vanishing convex combination exists. Now, small movements in enzyme space can improve an objective without compromising the others – a sign of a non-optimal state! In a Pareto-optimal state (lower left) this would be impossible.

In a metablic optimality problem, metabolic targets can be combined in an infinite number of ways (as convex combinations with different numerical weights), leading to an infinite number of single-objective extremality problems, each with a different objective function and a different solution. These solutions yield exactly the multi-objective-optimal points, i.e. the Pareto front of our original multi-objective problem. Moreover, with fluxes and metabolite concentrations as free variables, we obtain again the same solutions. Various formulations of the optimality problem lead to the same results!

## 8 Discussion

Cell fitness may depend on all kinds of things, including a fast replication of cell components or on complex dynamic behaviour. However, our objectives in pathway models are often much simpler: they are given by a function of fluxes, metabolite concentrations, and enzyme levels. How can we make a logical link between such pathway objectives and cell fitness? Even if we assume that fitness is a direct function of “biomass production’, this production may happen outside our pathway of interest, and our pathway would contribute indirectly (in the sense that without our pathway, biomass production would stop, or run more slowly). To quantify this indirect contribution, we define a (local) pathway objective as an “optimistic” proxy for cell fitness. An optimistic pathway objectives describes what fitness the cell can maximally achieve with a given state of the pathway. For simplicity, we further approximate that the pathway objective by a sum of targets for fluxes, metablite concentrations, and enzyme levels, in order to optimise or constrain these targets separately, or to use them for multi-objective optimisation.

We saw that metabolic optimality principles can be unified by describing them on the state manifold. A similar unification exists in classical thermodynamics, where variables such as temperature, pressure, and volume of a gas are related by a state equation, defining the possible states. These variables are interdependent, and there is no “natural choice” of basic variables: sometimes, it may be practical to choose volume and energy, or another time pressure and temperature as the basic system variables. In metabolic models, similarly we may parameterise metabolic states by enzyme levels or by metabolite concentrations and fluxes (see Figs (3) and (6)) – two ways to describe the state manifold! By picturing states on a state manifold (and not just, for example, in flux space), we acknowledge that nature does not distinguish between free and dependent variables, and that our choice of basic variables is nothing but a model assumption^24^. This description also shows how different modelling approaches (such as FBA, RBA, kinetic models, and coarse-grained cell models) are logically related and facilitates modular or layered modelling.

If the state manifold is parameterised by metabolite concentrations and fluxes, our general optimality problems can be written as optimality problems in these variables. For example, to maximise the fitness Eq. (1) at given fluxes, we can optimise a function of metabolite concentrations (knowing that enzyme levels depend uniquely on metabolite concentrations and fluxes). The resulting optimality problems have particular advantages and disadvantages. With enzyme levels as choice variables, it is easy to constrain them by a bound on the sum of enzyme levels. A layered flux/metabolite optimisation has different advantages. It can be split into separate layers a concave optimality problem for fluxes [18] and a convex optimality problem for metabolite concentrations [12] which are easier to study and solve. Moreover, our main metabolic target variables, the fluxes, can easily be predefined or constrained. This method can also help us define plausible flux cost functions *a*(**v**) for flux analysis. Linearised versions of these cost functions can be used directly in FBA, for instance in FBA with molecular crowding. With kinetic models being the “gold standard”, this justifies FBA as a method, first, by justifying flux cost functions and enzyme constraints in general, and second, by showing how they depend on the details of an underlying kinetic model (kinetic constants, thermodynamic forces, or external metabolite concentrations).

For any subsystem of a cell (and in fact, for any subset of variables), effective objectives can be defined as “optimistic objectives”. Enzymatic flux cost functions describe overhead costs: they formally appear as direct costs of a flux, but areis caused or implied indirectly in reality. The *enzymatic flux cost*, for example, describes an enzyme demand. The fact that they arise elsewhere (e.g. in enzyme production) is not shown explicitly in the formulae. When deriving metabolic optimality problems from an optimality problem on the state manifold, overhead costs arise as effective, optimistic costs due to hidden variables: if enzyme levels are hidden (i.e. projected out), their cost still remains as part of an optimistic (cost) objective for fluxes. Generally, overhead costs can describe two things: costs that arise from *causal conditions* (e.g. the cost of enzyme levels, as a condition for a flux) and costs that arise from *consequences or side-effects* of a process (e.g. the cost of a toxic side product). However, a strict distinction of this sort is difficult because in a cell, instead of a simple sequential causality, there are cycles and feedback loops. If we think of an entire cells, even enzyme cost itself is not a “basic” cost function, but an overhead cost: higher enzyme levels are not directly costly, but because of their costly effects on the cell state: protein production causes costs for precursors and ribosomes, and proteins consume precious space (for which they compete with other growth-relevant processes). This hodl for any cell variable: if we follow these chains of dependencies, we never find an “ultimately costly” variable: instead, we will go in cycles and see lots of constraints, and the enzyme costs that we first localised “at the boundary of our pathway” turn out to be an effective decription of opportunity costs in the entire (growing and living) cell. We saw that such opportunity costs can be approximated by optimistic costs with respect to our pathway, if optimality is assumed.

Whether a target function (such as enzyme cost or biomass production) is used as an objectives or to formulate a constraint depends on how we believe cells manage cellular variables – i.e. whether a target is for some reason is fixed, or whether it can be varied at some cost. This decision is not mainly about facts in reality, but rather a model assumption. For example, to model the compromise between biomass production and enzyme demand, we may fix the amount of enzyme to be allocated to metabolic reactions or pathways: in this case, the enzyme amount in each pathway can be increased at the expense of others. if we model one pathway, we may describe this negative impact on other, non-modelled pathways by an effective (overhead) enzyme cost. We can extend this argument to the whole metabolic network: if metabolic production is really important, cells will reallocate protein resources from other processes (e.g. ribosome production) to metabolism; and if some system is used and requires resources, the resulting loss in other systems can be described as an opportunity cost for our metabolic enzymes, to be used as a cost function. But the enzyme budget cannot be infinitely increased. Eventually, an optimal protein compromise between all cell functions will be reached, including metabolic enzymes, transporters, ribosomal proteins, chaperones, and so on. If this optimal state exists, we can take the metabolic enzyme budget *in this compromise state* as a *fixed* enzyme budget for metabolism. This brings us back to a “fixed-budget” model. Whether restrictions on cell variables should be described by objectives or constraints is therefore not a matter of physics or physiology, but a modelling decision: a choice between possible variation scenarios that we use to describe the possibilities of a cell. In our formulae, constraints can be replaced by cost terms and cost terms can be replaced by constraints, depending on what “freedoms” we attribute to the cell in our variation scenarios. Multi-objective optimisation, finally, can describe the full set of solutions that might be optimal for cells under different external conditions. Formally, each of these situations can be described by fixing some of the targets, giving rise to a single-objective optimality problem with specific weights or constraints for the metabolic targets.

If we study optimal metabolic behaviour with simple notions of optimality, what can we learn about actual cells? Metabolic economics does not claim optimality in reality, and optimality in cells can neither be proven nor disproven. Here, we take optimality as a model assumption and draw our conclusions. By asking how optimal behaviour, if it existed, *would* shape metabolic states, we can revisit some tacit assumptions made by biologists and study their consequences: using models, we can simulate how well organisms function and how selection pressures and constraints are expected to shape them. In [12], we tested directly the predictive power of an optimality principle, the principle of minimal enzyme cost. We considered a kinetic model with given flux distribution and predicted enzyme and metabolite concentrations in two ways: first, by sampling possible metabolite concentrations and solving for enzyme levels, and second, by optimising metabolite and enzyme levels for a minimal total enzyme demand. the second type of predictions was much better supported by data, which speaks in favour of our optimality principle. However, this test has not been repeated for other cases. In general, we cannot expect strict optimality in cells, and in fact they do not strictly save enzyme resources. Even if cells were evolutionarily optimised, they would still behave non-optimally under unpredictable experimental conditions or after “unusual” perturbations such as gene knockouts.

If cells in reality behave “non-optimally” and are subject to multiple, complicated requirements, then what is the point in considering optimality frameworks that favour extremely streamlined and specialised cell states? There are a set of good reasons. On the one hand, optimal states can be seen as “ideal” states that cannot be actually reached, but that see how much fitness fitness (e.g. productivity) is “lost” in real cell states. On the other hand, we may stick to the optimality assumption, but assume more realistic fitness objectives or constraints. For example, phenomena such as preemptive gene expression or bet hedging in cell subpopulations, leading to bacterial persistence, may be understood as adaptations to complex environments with varying supplies or with challenges by antibiotics. To describe such behaviours as beneficial, we may include adaptations to unexpected future challenges as side objectives into our optimality problems. Then, “wasteful” enzyme profiles turn out to be economical, e.g. as part of a bet-hedging strategy. But even if we don’t believe that cells, in reality, behave optimally, optimality considerations can be useful! By comparing real cell states to ideal optimal states (considering simple objectives), we may learn how clsoe cells are to optimal states, what prevents them from reaching these states, and what are plausible objectives to explain observed behaviour of cells.

The aim of this article – outlining a general theory of optimal metabolic states – has been achieved only partially. Optimality problems on metabolic networks can be formulated on the state manifold, that they can be solved by ECM or FCM, and we derived general optimality conditions. However, this framework is not ideally suited for modular modelling, for models that allow us to “zoom in and out”, or for detailed models of entire cells. First, it lacks some important features, including density constraints for metabolites and proteins (in different cell compartments), dilution fluxes, and non-enzymatic reactions. Second, in a pathway model, the optimal state depends on variables *inside* the pathway, not on (unknown) variables in the surrounding network. However, the optimality conditions for an enzyme level do depend on effects anywhere in the network, also outside our pathway model! A unified theory should be invariant against “zooming”: if we model a network region, our optimality principle can must formulated entirely in terms of variables in this region! But in reality, if we model a single pathway, cell fitness will always depend on processes outside the pathway. But how can we make a theory “zoomable”, i.e. applicable to any part of the cellular networks? Often none of the pathway variables will directly appear in the cell fitness function, and the control coefficients between pathway variables and the actual fitness-relevant variables amy be unknown! How can we solve this paradox? Of course, we may predefine pathway objectives at will, regard them as proxies for cell fitness, and hope for the best – but this would be tinkering rather than developing a consistent theory.

A consistent economic theory of metabolism should not only hold for a single model, but should also allow us to move flexibly between models in order to to embed pathway models into larger networks, to derive pathway objectives from a pathway’s effects on cell fitness, and to describe models on different scales of resolution in a single, consistent framework. It should be general enought to describe local (pathway) and global (cell) models by one formalism. For predictions it should not matter whether metabolic targets are bounded or optimised, because this is a question of model formulation, not a fact about reality. Finally, the theory should be local, i.e. applicable to any region of the cellular network without requiring knowledge about the rest of the network. But is this possible? If we zoom into a network region, we lose all information about the surrounding system! So how can we deal with objectives that depend on the outer variables? In [21], I sketch a theory that has all these features: a theory in which metabolites, enzymes, and reactions carry economic values. These value variables are dual to our physical model variables (concentration, fluxes, or production rate) and describe how variations of our physical variables affect fitness, just like forces are dual to displacements in mechanics, pressure is dual to volume in thermodynamics, and prices are dual to amounts in economics. Using these economic values, we can define meaningful local pathway objectives that represent a pathway’s contribution to cell fitness, which makes the theory local. The new formalism can also handle new details such as bounds on protein budgets, metabolite dilution, or non-enzymatic reactions, which makes it widely applicable to metabolic models and cell models.

## Abbreviations

ECM: Enzyme cost minimisation
FCM: flux cost minimisation

## Acknowledgements

I thank Mariapaola Gritti, Elad Noor, and Stefan Müller for thinking with me. This work was funded by the German Research Foundation (Ll 1676/2-2).

## A Probability distributions on the state manifold

Aside from objective functions, probability distributions are another important type of functions on the state manifold. By definition, probability density function must be non-negative function and must have an integral of 1 on the state manifold (or on our region of interest). By integrating the probability density function over a region of the manifold, we obtain the finite probabilities for the system state to be in this region. Probability distributions can describe states in a cell population or our (uncertain) knowledge about the state of specific cells. In practice, probability distributions can also be used for a sampling of cell states.

Probability densities can be directly derived from objective functions, assuming that profitable states are also in fact more probable. The idea comes from physics. A probability density can always be written in the form 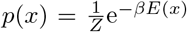, using a real-valued energy function *E*(*x*), where *β* is a scaling parameter for *E* and *Z* (the “state sum”) is chosen to scale the integral to 1. In statistical physics, *E*(*x*) is the energy of a state, *β* represents an inverse temperature, and *p*(*x*) is the Boltzmann probability density. Here we can use the same formula, but with a different interpretation. By treating the negative fitness (e.g. a cost or a negative benefit) as an “energy function” *E*, we can define probability distributions over metabolic states analogous to the Boltzmann distribution in statistical mechanics. In this distribution, the probability states increases with their fitness. The “strictness of selection”, which translates fitness into probabilities can be tuned by a parameter *β*, corresponding to an inverse temperature. In the limit of infinite strictness (“zero temperature”), all non-optimal states have zero probability. In the limit of zero strictness (“infinite temperature”), we obtain a uniform distribution, with all states being equally likely. “Temperatures” in between lead to distributions in between, with higher probabilities for high-fitness states.

## Footnotes

2 In this paper, “pathway” may denote any part of a metabolic network or even the network as a whole. The term “pathway” emphasises that any metabolic system is surrounded by even larger networks.

3 If we describe fitness by a simple difference of benefit and cost terms (for fluxes and concentrations) this is just for convenience. For the most part, the theory holds for general fitness functions ℱ(**v, c, e**) if they are reasonably well behaved (in particular, differentiable) and increase monotonically with the enzyme levels. Instead of three terms as in Eq. (1), we may also use formulae such as ℱ(**v, c, e**) = *q*(**v, c**) − *h*(**e**) (“metabolic objective minus enzyme cost”) or ℱ(**v, c, e**) = *b*(**v**) − *h*^kin^(**c, e**) (“flux benefit minus kinetic cost”). A practical use of such additive functions is to keep some terms fixed while optimising others. But we may split our objective even further into multiple sub-objectives, e.g. to score the production of different compounds by separate benefit terms. Trade-offs between such objectives can be treated like the trade-offs between flux and metabolite objectives.

4 Which metabolites are described as external (concentrations given by model parameters) or internal (concentrations described by variables) depends on the modeller’s choice.

5 To obtain a unique mapping between reactions and enzymes, we can formally introduce “monoreactions” (a reaction catalysed by different isoenzymes is described by separate monoreactions) and “monoenzymes” (a multifunctional enzyme, catalysing several reactions, is described by separate monoenzyme pools).

6 If we put a positive lower bound on |∂*a/*∂*v*_*l*_|, the flux cost must show a kink^7^ at *v*_*l*_ = 0. For flux cost functions representing enzyme cost in kinetic models [25, 18], we can show that (∂*a/*∂*v*_*l*_) *v*_*l*_ = (∂*h/*∂*e*_*l*_) *e*_*l*_ *>* 0).

8 Considering density constraints Σ_*i*_ *α*_*i*_ *c*_*i*_ *≤ d*_tot_ leads to convex upper bound in log concentration space, cutting the polytope (the resulting set is not a polytope anymore, but convex, so minimisation problems in this space remain tractable).

9 High concentrations may cause problems because of limited space in cells (in a densely packed cell, the osmotic pressure is high and diffusion is slow, which limits enzymatic rates) or because of toxicity. Low concentrations can make reactions kinetically inefficient (if this concerns reactions outside the model, the resulting loss can be captured by a metabolite cost term); at very low concentrations, below an average molecule number of 1 per cell, metabolites are effectively absent.

10 By definition, an algebraic manifold 𝒦 is the solution set of a system of polynomial equations. This holds for our state manifold: (i) Σ_*l*_ *n*_*il*_ *v*_*l*_ = 0 is linear and therefore a polynomial in the *v*_*l*_; (ii) the rate laws have the form *v*_*l*_ = *a*_*l*_(*e*_*l*_, **c**)*/b*_*l*_(**c**), with polynomials *a*_*l*_ and *b*_*l*_. We can rewrite this as *v*_*l*_ *b*_*l*_(**c**) − *a*_*l*_(*e*_*l*_, **c**) = 0, which is a polynomial in the variables *v*_*l*_, *c*_*i*_, and *e*_*l*_.

11 Compared to a single reaction, we encounter some complications: (i) Instead of a single flux direction, we need to consider flux patterns describing all flux directions in the network. Each feasible flux pattern defines a feasible F-polytope in flux space and a feasible M-polytope in metabolite space. (ii) Internal metabolites must be mass-balanced, to ensure stationary fluxes. (iii) Flux profiles may be constrained to a predefined flux benefit. Because of the conditions (ii) and (iii), some patches will be discarded. (iv) The enzyme demand function **e**(**v, c**) is multidimensional. (v) Concentration ranges or fixed concentrations may be defined separately for each metabolite.

12 To show this, we imagine that a cell, starting in an initial state, first decreases all enzyme levels and fluxes gradually to zero, then gradually changes the metabolite profile, and finally increases fluxes and enzyme levels to get to the end state. However, this path leads through a nonviable (thermodynamic equilibrium) state in which all metabolic fluxes vanish. Possibly, the viable states are not path-connected. Moreover, if we restrict the state manifold to some positive flux benefit, the proof will not apply anymore. In reality, on their way between steady states, cells may have to pass through non-steady or unsustainable states.

13 If several of these steady-state solutions are dynamically stable, we call this multi-stability.

14 If we use thermodynamically consistent, reversible rate laws, the enzyme demand **e**(**v, c**) is positive everywhere on the set of thermodynmically feasible states (**v, c**).

15 There is one catch: as mentioned above, the functions **c**^st^(**e**) and **v**^st^(**e**) (even at predefined conserved moiety concentrations) may be non-unique (in cases of multi-stability). This means for describing steady states by functions in e-space, we additionally need to require that we stay on the same sheet (in (**v, c, e**) space)

17 For models without conserved moieties, the proof of Eq. (18) is relatively simple. We start from an optimal state and apply a metabolite variation *δ***c**, together with a flux-compensating enzyme variation *δ***e** satisfying 0 = *δ***v** = **E**_c_ *δ***c** + **E**_e_ *δ***e**, hence given by 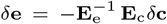. Since the original state was already optimal, the cost of this compensated variation must vanish: 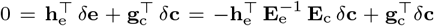. This yields 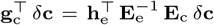. Because the same logiv must hold for any variation *δ***c**, we obtain Eq. (18).

18 Here we consider “infinitely small” variations which capture first-order effects and are mathematically defined by differential forms.

19 The fact that metabolite price and adjacent enzyme prices are balanced can be seen as an example of the “willingness to pay” principle: decreasing the metabolite concentration (and thus the metabolite cost) would be possible, but the cell would have to “pay” an exactly equal cost (for adjusted enzyme levels).

20 The same logic also holds for general fitness functions *f* (**v, c, e**) as long as the enzyme prices −∂*f/*∂*e* are positive.

21 We can define such variations by setting *δ***c**_‖_ = **L** *δ***c**^ind^ and *δ***v**_‖_ = **K** *δ***v**^ind^, where **c**^ind^ and **v**^ind^ denote vectors of independent metabolite concentrations and fluxes. If we insert this into Eq. (22), we obtain laws for variations **c**^ind^ and **v**^ind^ in which non-locality (due to the matrices **L** and **K**) becomes obvious.

22 For simplicity, we consider linear combinations *B* = Σ_*i*_ *β*_*i*_ *b*_*i*_. Nonlinear combinations would lead to optimality conditions of the same form, but with variable prefactors *β*_*i*_ in the balance of gradients are not fixed. They are not constant parameters, but given by derivatives ∂*B/*∂*b*_*i*_ of the combined objective with respect to the single objectives, in the metabolic state in question.

23 To find an optimal metabolic state, constrained optimisation can be applied in different ways: a *nested* optimisation of several objectives results in layers of restricted optimisation, while the overall solution does not depend on the order of nesting. In contrast, in a *series* of optimisations we first optimise objective A; then we optimise B, while restricting A to its optimal value, and so on. Here the order of optimisations plays a role. This procedure is used in the principle of minimal fluxes.

24 Ideally, reformulating a model with a different choice of basic variables should not change the results. In practice, however, there may be differences because of tacit assumptions, for instance, about probability distributions: for example, a uniform sampling of fluxes (which determine optimal enzyme and metabolite profiles) will not yield the same results as a uniform sampling of enzyme profiles (which determine metabolite and flux profiles).

